# ZBP1 induces inflammatory signaling via RIPK3 and promotes SARS-CoV-2-induced cytokine expression

**DOI:** 10.1101/2021.10.01.462460

**Authors:** Ruoshi Peng, Xuan Wang-Kan, Manja Idorn, Felix Y Zhou, Susana L Orozco, Julia McCarthy, Carol S Leung, Xin Lu, Katrin Bagola, Jan Rehwinkel, Andrew Oberst, Jonathan Maelfait, Søren R Paludan, Mads Gyrd-Hansen

## Abstract

COVID-19 caused by the SARS-CoV-2 virus remains a threat to global health. The disease severity is mediated by cell death and inflammation, which regulate both the antiviral and the pathological innate immune responses. ZBP1, an interferon-induced cytosolic nucleic acid sensor, facilitates antiviral responses via RIPK3. Although ZBP1-mediated cell death is widely described, whether and how it promotes inflammatory signaling is unclear. Here, we report a ZBP1-induced inflammatory signaling pathway that depends on ubiquitination and RIPK3’s scaffolding ability independently of cell death. In human cells, ZBP1 associates with RIPK1 and RIPK3 as well as ubiquitin ligases cIAP1 and LUBAC. RIPK1 and ZBP1 are ubiquitinated to promote TAK1- and IKK-mediated inflammatory signaling. Additionally, RIPK1 recruits the p43/41-caspase-8-p43-FLIP heterodimer to suppress RIPK3 kinase activity, which otherwise promotes inflammatory signaling in a kinase activity-dependent manner. Lastly, we show that ZBP1 contributes to SARS-CoV-2-induced cytokine production. Taken together, we describe a ZBP1-RIPK1-RIPK3-mediated inflammatory signaling pathway relayed by the scaffolding role of RIPKs and regulated by caspase-8. Our results suggest the ZBP1 pathway contributes to inflammation in response to SARS-CoV-2 infection.

## Introduction

Viruses can cause a range of severe acute and chronic diseases and represent a global health threat, as illustrated by Severe Acute Respiratory Syndrome Coronavirus 2 (SARS-CoV-2) that caused the Coronavirus disease (COVID-19) pandemic. Many viral diseases are characterized by over-reaction of the immune system that contributes to disease development (Fajgenbaum & June, 2020). This is seen in COVID-19 where patients with severe symptoms suffer from a “cytokine storm”, which is featured by elevated excessive circulating cytokine levels that correlate with lethality (Zhou *et al*, 2020; Chen *et al*, 2020; Lucas *et al*, 2020). Inflammation and cell death underlie antiviral innate immune responses and contribute to pathological inflammatory conditions when deregulated. Receptor interacting protein (RIP) kinases (RIPKs), engaged downstream of immune receptors, are central regulators of cell death and inflammatory signaling pathways and contribute to host immune defenses against viruses and bacteria (Newton, 2020; He & Wang, 2018; Topal & Gyrd-Hansen, 2021).

Z-DNA-binding protein 1 (ZBP1) is a cytosolic nucleic acid sensor and an interferon-induced pattern recognition receptor (PRR) important for antiviral immune responses (Kuriakose & Kanneganti, 2018; Pham *et al*, 2013; Omoto *et al*, 2015; Upton *et al*, 2012; Kuriakose *et al*, 2016; Daniels *et al*, 2018; Kesavardhana *et al*, 2017; Maelfait *et al*, 2017; Thapa *et al*, 2016; Takaoka *et al*, 2007). After activation, ZBP1 recruits RIPK3 and RIPK1 to execute programmed cell death, including apoptosis, necroptosis, pyroptosis or a mixture of them, depending on the cell type and caspase activity (Upton *et al*, 2012; Thapa *et al*, 2016).

RIPK3 mediates necroptosis by phosphorylating MLKL, which in turn oligomerizes and forms pores in the cell membrane (He *et al*, 2009; Cho *et al*, 2009; Sun *et al*, 2012). In addition to its kinase domain, RIPK3 contains a RIP Homotypic Interaction Motif (RHIM) that mediates its recruitment to other RHIM-containing proteins, namely RIPK1, the Toll-like receptor (TLR) adaptor TIR-domain-containing adapter-inducing interferon-β (TRIF) and ZBP1 (He *et al*, 2009; Kaiser *et al*, 2013; Rebsamen *et al*, 2009). The activation of RIPK3 is proposed to occur within a RHIM-mediated oligomer enucleated by the RHIM of RIPK1 and stabilised by phosphorylation of RIPK1 and RIPK3 molecules (Li *et al*, 2012; Wu *et al*, 2014). In addition to its necroptosis-promoting activity, RIPK3 has been suggested to promote inflammatory signaling during TNF- and TLR-induced necroptosis and downstream of ZBP1 (Zhu *et al*, 2018; Muendlein *et al*, 2020; Najjar *et al*, 2016; Rebsamen *et al*, 2009). However, the mechanism of RIPK3-mediated inflammatory signaling remains unresolved.

The formation of non-degradative ubiquitin (Ub) chains linked via lysine 63 (K63-Ub) and methionine 1 (M1-Ub) within receptor signaling complexes facilitates the activation of the kinases TAK1 and IKKα/β, which in turn activate MAP kinase signaling and NF-κB signaling to stimulate the expression of pro-inflammatory cytokines and chemokines (reviewed in Hrdinka and Gyrd-Hansen, 2017).

In this study, we identify RIPK1 and RIPK3 as scaffolding kinases that mediate ZBP1-triggered inflammatory signaling independently of cell death. ZBP1-RIPK3-RIPK1 inflammatory signaling is dependent on K63-Ub and M1-Ub and Ub ligases cIAPs and LUBAC but does not require the kinase activity of RIPK1 and RIPK3. Inhibition of caspase-8 activity exposes a RIPK3 kinase activity-mediated inflammatory signaling pathway, akin to the pathway triggered by TNF when cIAPs and caspase-8 are inhibited. Finally, we provide evidence that ZBP1 contributes to the production of cytokines and chemokines during SARS-CoV-2 infection.

## Results

### ZBP1 and RIPK3 stimulate inflammatory signaling independently of cell death

ZBP1-induced signaling is mediated by RIPK3 and is dependent on RHIM interactions (Kaiser *et al*, 2008; Rebsamen *et al*, 2009). To investigate the ability of ZBP1 to stimulate inflammatory signaling versus cell death, we generated HT29 cells with doxycycline (Dox)-inducible expression of FLAG-tagged wild-type (WT) ZBP1 (ZBP1^WT^) or a mutant form in its Z-form nucleic acid binding (Zα)–domains (ZBP1^Zα1α2mut^) (Figure 1A). Dox treatment induced the expression of ZBP1^WT^ and ZBP1^Zα1α2mut^ in a dose-dependent manner, albeit ZBP1^Zα1α2mut^ expressed at higher levels than ZBP1^WT^ (Figure 1B).

**Figure 1.**
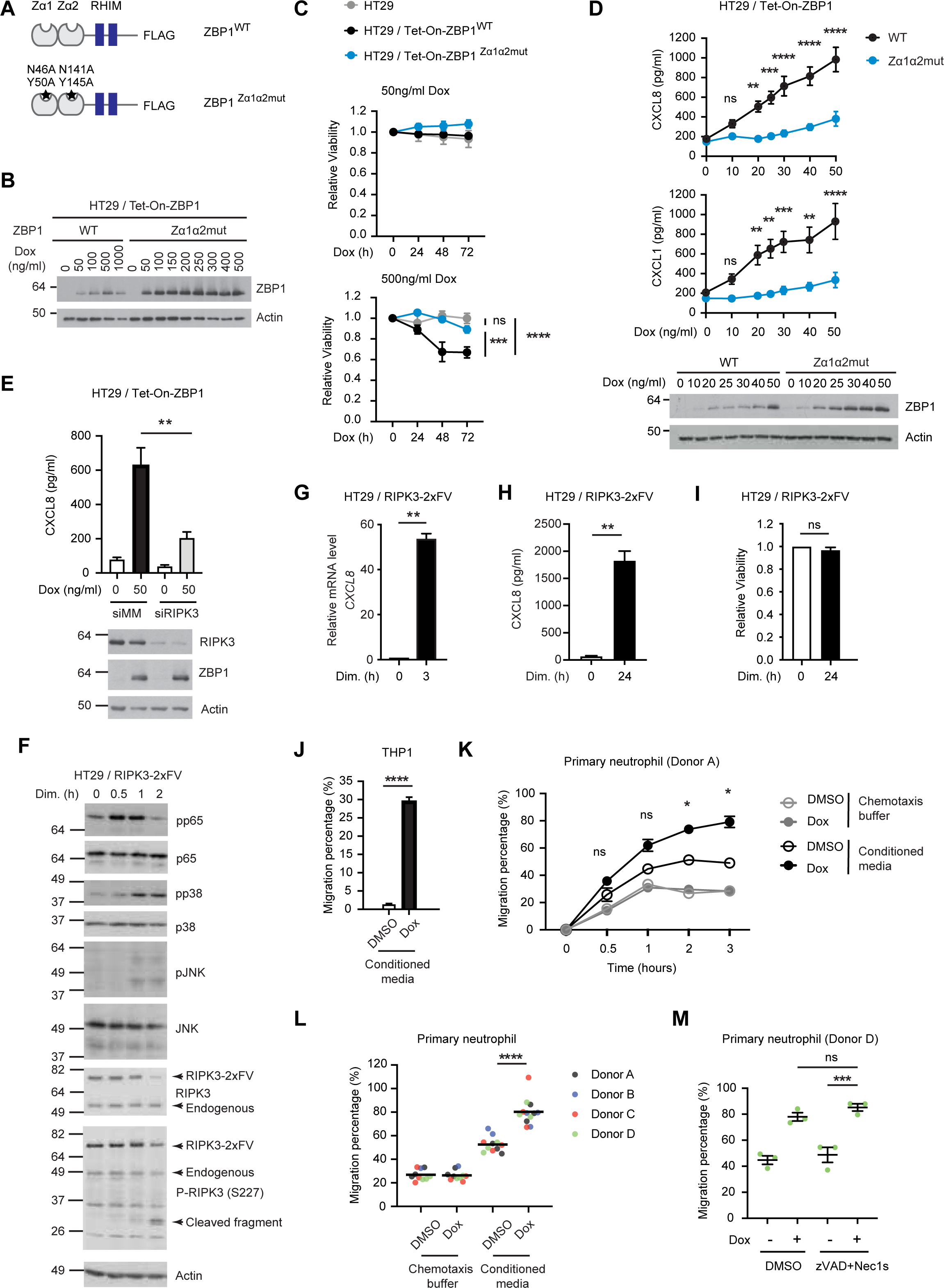
**See also Figures S1-S3. ZBP1 and RIPK3 stimulates inflammatory signaling independently of cell death** (A) Schematic illustration of WT and Zα1α2-mutant (Zα1α2mut) human ZBP1 induced to express in HT29 cells. (B) Western blot analysis of the dose-dependent expression of ZBP1 in HT29 / Tet-On-ZBP1^WT^ and HT29 / Tet-On-ZBP1^Zα1α2mut^ cells treated with indicated concentrations of Dox for 24 h. (C) Relative viability of indicated cells treated with Dox for 72, 48, 24 or 0 h. Cell viability at the end of treatment was determined by measuring ATP levels. Measurement values were normalized to that of 0 h for each cell line. Data is presented as mean with S.E.M (n = 3). Two-way ANOVA and Tukey’s multiple comparisons tests were used to test for the statistical differences between different cell lines. ns (not significant), p = 0.9510, ***, p = 0.0001, ****, p < 0.0001. (D) Chemokine concentrations in cell culture supernatants of HT29 cells induced to express ZBP1^WT^ or ZBP^Zα1α2mut^ for 24 h with indicated Dox concentrations. Data is presented as mean with S.E.M. (n = 4). Two-way ANOVA and Sidak’s multiple comparisons t-test were used to test for the statistical differences between the two cell lines at each concentration. ns, p > 0.5, **, p = 0.0035 for CXCL8 at 20 ng/ml, p = 0.0067 for CXCL1 at 20 ng/ml, p = 0.0020 at 50 ng/ml, p = 0.0013 at 40 ng/ml, ***, p = 0.0004 for CXCL8 at 25 ng/ml, p = 0.0008 at 30 ng/ml, ***, p < 0.0001. Cells from the same wells were lysed and analyzed by western blotting. (E) CXCL8 concentrations in cell culture supernatants of HT29 / Tet-On-ZBP1 cells transfected with siRNA targeting mismatch sequence (MM) or RIPK3 and treated with Dox at indicated concentrations for 24 h. Data is presented as mean with S.E.M. (n = 4). Unpaired t-tests were used to test for statistical differences between indicated conditions. **, p = 0.0079. Cells were lysed and analyzed by western blotting. (F) Western blot analyses of inflammatory marker proteins in HT29 / RIPK3-2xFV cells treated with 100 nM dimerizer. Arrows indicate RIPK3 signals. (G) Relative *CXCL8* mRNA levels in HT29 / RIPK3-2xFV cells treated with 0 or 100 nM dimerizer for 3 h. Data is presented as mean with S.E.M. (n = 3). A Welch’s t-test was used to test for statistical differences between indicated conditions. **, p = 0.0016. (H) CXCL8 concentrations in supernatants of HT29 / RIPK3-2xFV cells treated with 0 or 100 nM dimerizer for 24 h. Data is presented as mean with S.E.M. (n = 3). A Welch’s t-test was used to test for statistical differences between indicated conditions. **, p = 0.0093. (I) Relative viability of HT29 / RIPK3-2xFV cells treated with 0 or 100 nM dimerizer in the presence of 0.5 µg/ml mouse IgG. Data is presented as mean with S.E.M. (n = 3). A Welch’s t-test was used to test for statistical differences between indicated conditions. ns, p = 0.3268. (J) Transwell migration percentages of THP1 cells towards conditioned media from HT29 / Tet-On-ZBP1 cells treated with DMSO or 500 ng/ml Dox for 24 hours. Data is presented as mean with S.E.M. (n = 3). An unpaired t-test was used to test for statistical differences between indicated conditions. ****, p < 0.0001. (K) Transwell migration percentage of primary neutrophils from Donor A towards conditioned media from HT29 / Tet-On-ZBP1 cells treated with DMSO or 500 ng/ml Dox for 24 hours, or chemotaxis buffer containing equal volume and amount of DMSO or Dox. Data is presented as mean with S.E.M. (n = 3 biological replicates of conditioned media). Two-way ANOVA and Dunnet’s multiple comparison tests were used to test for statistical differences between the migration percentage of neutrophils towards conditioned media from DMSO-treated cells and that from Dox-treated cells at each time point. ns, p = 0.3110 for the 0.5 h time point, p = 0.0861 for the 1h time point. *, p = 0.0113 for the 2 h time point, p = 0.0133 for the 3 h time point. (L) Transwell migration percentages of primary neutrophils towards conditioned media from HT29 / Tet-On-ZBP1 cells treated with DMSO or 500 ng/ml Dox for 24 hours, or chemotaxis buffer containing equal volume and amount of DMSO or Dox. Data is presented as individual values with grand mean (n = 10 of chemotaxis buffer containing DMSO or Dox, n = 11 of conditioned media from DMSO-treated cells, n = 12 of conditioned media from Dox-treated cells). An unpaired t-test was used to test for statistical differences between indicated conditions. ****, p < 0.0001. (M) Transwell migration percentage of primary neutrophils towards conditioned media from HT29 / Tet-On-ZBP1 cells treated with 0 or 500 ng/ml Dox in combination with DMSO or 20 µM zVAD + 10 μM Nec1s for 24 h. Data is presented as individual values with mean and S.E.M. (n = 3). One-way ANOVA and Sidak’s multiple comparisons test were used to test for statistical differences between indicated conditions. ns, p = 0.4129, ***, p = 0.0004.

In accordance with the ability of ZBP1 to stimulate ligand-dependent cell death (Jiao *et al*, 2020; Wang *et al*, 2020), treatment with 500 ng/ml Dox resulted in the loss of viability after 48 h in HT29 cells expressing ZBP1^WT^ but not ZBP1^Zα1α2mut^ (Figure 1C). Contrary to previous studies in murine systems (Upton *et al*, 2012; Wang *et al*, 2020; Zhang *et al*, 2020; Jiao *et al*, 2020), the cell death induced by ZBP1 in HT29 cells was predominantly apoptosis, as viability was restored by the caspase inhibitor zVAD-fmk (zVAD) alone or in combination with inhibition of RIPK3 (GSK’872), RIPK1 (Nec1s) or MLKL (necrosulfonamide; NSA), whereas GSK’872, Nec1s or NSA alone did not restore viability (Figure S1A).

Treatment with 50 ng/ml or 100 ng/ml Dox did not result in the loss of viability although ZBP1 expression was induced (Figures 1C and S1B). Based on this, we treated the cells with increasing concentrations of Dox up to 50 ng/ml and measured cytokine production after 24 h. Despite no loss of cell viability, the induced expression of ZBP1 stimulated the production of chemokines CXCL8 and CXCL1 in a Dox concentration-dependent manner, which was also dependent on the ligand-binding ability of ZBP1 (Figure 1D). The Dox-induced expression of ZBP1^Zα1α2mut^ was higher than ZBP1^WT^, yet chemokine production was significantly lower, underscoring the importance of ligand-binding for ZBP1 signaling at low expression levels (Figure 1D). Similarly, transient expression of ZBP1 in HEK293T cells induced NF-κB activity in a ligand-dependent manner (Figure S1C). In line with previous reports (Rebsamen *et al*, 2009; Kaiser *et al*, 2008), ZBP1-induced inflammatory signaling relied on RIPK3’s RHIM, since ZBP1^WT^ induced NF-κB activity in cells expressing full-length RIPK3 (RIPK3^FL^) but not a C-terminal truncated RIPK3 lacking the RHIM (RIPK3^dC^) (Figure S1C). Similarly, ZBP1-induced inflammatory signaling in HT29 cells was mediated by RIPK3 (Figures 1E and S1D). Notably, at higher Dox concentrations, both ZBP1^WT^ and ZBP1^Zα1α2mut^ induced the production of chemokines in HT29 cells, either when treated with equal amount of Dox (Figure S1E) or when induced to express at the same level (Figure S1F), albeit the chemokine production induced by ZBP1^Zα1α2mut^ was significantly lower than that by ZBP1^WT^ when expressing at the same levels (Figure S1F). This suggests that ZBP1, when highly expressed, can trigger inflammatory signaling independently of ligand-binding (Maelfait *et al*, 2017).

As RIPK3 is required for ZBP1-induced inflammatory signaling, we sought to determine if human RIPK3, like ZBP1, could stimulate inflammatory signaling without concomitant stimulation of cell death. We employed the homodimerization domain B (DmrB, FK506-binding protein F36V mutant, hereafter FV)-mediated RIPK3 oligomerization system, which allows for acute and controlled activation of RIPK3 by B/B homodimerizer AP20187 (hereafter referred to as dimerizer) (Clackson *et al*, 1998; Orozco *et al*, 2014; Yatim *et al*, 2015; Rodriguez *et al*, 2016). Chimeric RIPK3 fused with FV domains (RIPK3-1xFV, RIPK3-2xFV and RIPK3dC-2xFV) was stably expressed in HT29, HCT116, U2OS and HEK293T cells (Figure S2A). Dimerizer-induced RIPK3 oligomerization in HT29 cells stimulated inflammatory signaling after 30-60 min as determined by phosphorylation of the NF-κB subunit RelA/p65 and MAPKs p38 and JNK as well as the expression and production of the chemokine *CXCL8* (Figures 1F-1H). We obtained similar results after dimerization or oligomerization of RIPK3 in HCT116, U2OS and HEK293T cells (Figures S2B-H). Notably, contrary to murine cells where oligomerization of RIPK3 induces both necroptosis and inflammatory signaling (Orozco *et al*, 2014; Yatim *et al*, 2015), RIPK3 oligomerization did not result in loss of viability of HT29 or HCT116 cells within 24 h (Figures 1I and S2I) although the HT29 cells were sensitive to TNF- induced cell death when pre-treated with Smac mimetic Compound A (CpA; (Vince *et al*, 2007)) (apoptosis) or CpA + zVAD (necroptosis) (Figures S2J).

In addition to CXCL8 and CXCL1, we found that ZBP1 expression in HT29 cells stimulated the secretion of the chemokines CXCL10/IP-10, CCL20, CXCL7 and other proinflammatory mediators (Figures S3A and S3B). This suggests that ZBP1 signaling may stimulate the chemoattraction of neutrophils and other immune cells. Indeed, conditioned media from Dox-treated HT29 / Tet-On-ZBP1 cells stimulated chemotactic migration of THP1 monocytic cells, neutrophil-like differentiated HL60 cells and primary human donor neutrophils (Figures 1J-1L and S3C). To determine if undetected cell death after Dox treatment might contribute to chemotaxis, HT29 cells were treated with zVAD and Nec1s in combination with Dox for chemotaxis. ZBP1-dependent chemotactic migration of primary donor neutrophils was unaffected by treatment with both inhibitors, indicating that the migration was stimulated by ZBP1-dependent inflammatory signaling and not by cell death (Figure 1M).

### ZBP1-RIPK3 inflammatory signaling is mediated by IKK and TAK1 independently of the kinase activity of RIPK3 or RIPK1

To gain insights into the mechanism underpinning ZBP1 and RIPK3-induced inflammatory signaling, HT29 cells were treated with kinase inhibitors in combination with Dox-induced expression of ZBP1. Treatment with Nec1s, GSK’872 and NSA did not lead to a significant decrease in ZBP1-induced production of CXCL8 and CXCL1 in HT29 cells or NF-κB activation in RIPK3-expressing HEK293FT cells (Figures 2A and 2B), indicating that inflammatory signaling by ZBP1 occurs independently of the kinase activity of RIPK3 and RIPK1. Accordingly, inhibition of RIPK3, RIPK1 or MLKL had no effect on the ZBP1-stimulated chemotactic migration of neutrophils (Figure 2C).

**Figure 2.**
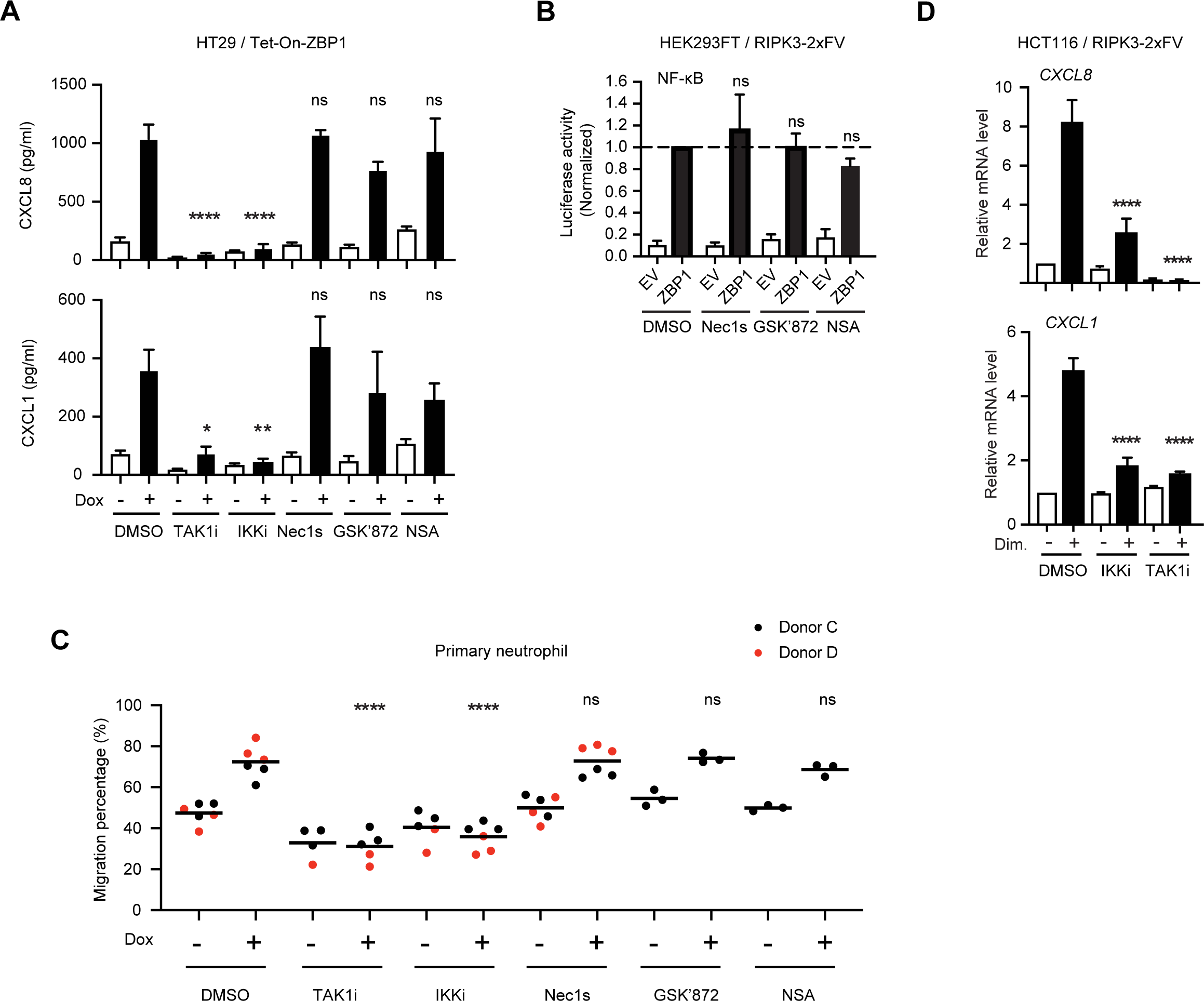
**ZBP1-RIPK3 inflammatory signaling is mediated by IKK and TAK1 but does not require RIPK3 and RIPK1 kinase activity** (A) Chemokine concentrations in cell culture supernatants of HT29 / Tet-On-ZBP1 cells treated with 0 or 50 ng/ml Dox in combination with indicated inhibitors for 24 h. TAK1i, 1 μM 5z-7-oxozeaenol. IKKi, 1 μM IKK Inhibitor VII + 5 μM IKK Inhibitor XII. Nec1s, 10 μM. GSK’872, 10 μM. NSA, 1 μM. Data is presented as mean with S.E.M. (n = 3). One-way ANOVA and Sidak’s multiple comparisons tests were used to test for statistical differences between Dox-induced samples pretreated with DMSO and corresponding inhibitor. ns, p > 0.2, *, p = 0.0104, **, p = 0.0041, ****, p < 0.0001. (B) Relative NF-κB activity in response to ZBP1 expression in the presence of indicated inhibitors. Cells were treated with indicated compounds immediately after transfection with dual luciferase plasmids and empty vector (EV) or ZBP1-expressing plasmids. Nec1s, 10 μM. GSK’872, 10 μM. NSA, 1 μM. Data is presented as mean with S.E.M. (n = 3). One-way ANOVA and Sidak’s multiple comparisons tests were used to test for statistical differences between ZBP1-transfected conditions treated with DMSO and treated with indicated inhibitors. ns, p = 0.9994 for Nec1s, p > 0.9999 for GSK’872, p = 0.9358 for NSA. (C) Transwell migration percentages of primary neutrophils towards conditioned media from HT29 / Tet-On-ZBP1 cells treated with 0 or 500 ng/ml Dox in combination with indicated inhibitors for 24 h. Data is presented as individual values with grand mean. Two-way ANOVA and Sidak’s multiple comparisons test were used to test for statistical differences between indicated conditions. ****, p < 0.0001, ns, p > 0.9999 for Nec1s and GSK’872, p = 0.9645 for NSA. (D) Relative mRNA levels of *CXCL8* and *CXCL1* in HCT116 / RIPK3-2xFV cells pretreated with DMSO, TAK1 inhibitor, or IKK inhibitors for 1 h before treated with 0 or 100 nM dimerizer for 3 h. Data is presented as mean with S.E.M. (n = 3). One-way ANOVA and Sidak’s multiple comparisons tests were used to test for statistical differences between indicated conditions and DMSO+dimerizer–treated condition. ****, p < 0.0001.

Conversely, ZBP1-induced chemokine production and neutrophil migration were blocked by the TAK1 inhibitor 5z-7-oxozeaenol (TAK1i) (Wu *et al*, 2013) and IKK inhibitors (IKKi) (Waelchli *et al*, 2006; Christopher *et al*, 2007) (Figures 2A and 2C). TAK1 and IKK inhibition also antagonized RIPK3 oligomerization-induced expression of *CXCL1* and *CXCL8* (Figure 2D). Together, this showed that ZBP1-induced inflammatory signaling is mediated by MAPK- and NF-κB pathways and relies on RIPK3 and RIPK1 as scaffolds rather than their kinase activity.

### K63-Ub and M1-Ub facilitates ZBP1 inflammatory signaling

RIP kinases function as scaffolds by serving as substrates for non-degradative ubiquitination in inflammatory signaling (Ea *et al*, 2006; Hrdinka *et al*, 2018; Hasegawa *et al*, 2008; Damgaard *et al*, 2012), which prompted us to investigate ubiquitination events after ZBP1 induction. FLAG-tagged ZBP1 was immunoprecipitated from Dox-treated HT29 cells. As expected, RIPK3 and RIPK1 were both co-purified with ZBP1 (Figure 3A) (Rebsamen *et al*, 2009; Kaiser *et al*, 2008). Interestingly, ZBP1 also co-purified high molecular weight (MW) Ub-conjugates, confirmed by treatment of the immunoprecipitated material with the deubiquitinase USP21 (Figure 3A). Enrichment of endogenous Ub-conjugates by GST-1xUBA (Fiil *et al*, 2013; Hrdinka *et al*, 2016) revealed that ZBP1 expression increased the ubiquitination of RIPK1 and of ZBP1 itself, whereas ubiquitination of RIPK3 was not detected under these conditions (Figure 3B). Further, enrichment of K63- and M1-Ub by linkage-selective Ub binders (SUBs) (Fiil *et al*, 2013; Hrdinka *et al*, 2016) showed that RIPK1 and ZBP1 were both modified by K63-Ub, whereas M1-Ub appeared to predominantly accumulate on ZBP1 (Figures 3C and S4A).

**Figure 3.**
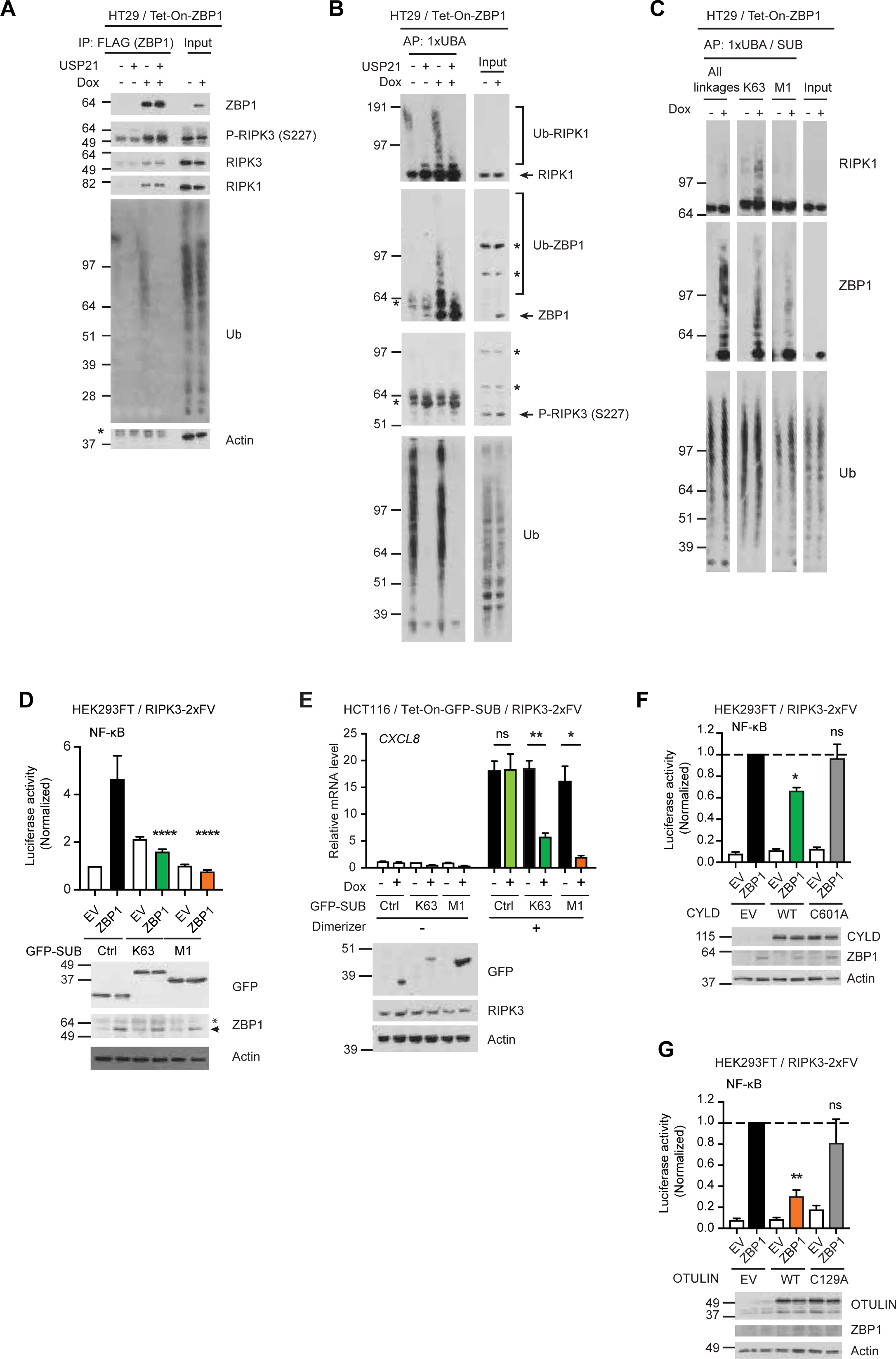
**See also Figure S4. K63-Ub and M1-Ub facilitate ZBP1 inflammatory signaling** (A) Western blot analyses of anti-FLAG (ZBP1) immunoprecipitation from HT29 / Tet-On-ZBP1 cells treated with 0 or 500 ng/ml Dox for 16 h. After enrichment, beads were incubated with 0 or 1 µM USP21 at 30°C for 1h for deubiquitination. Input loaded was 5% for ZBP1 and 1% for co-immunoprecipitants. Asterisk indicates antibody heavy chain signal in PD. (B) Enrichment of Ub-conjugates by GST-1xUBA for analysis of ubiquitination status of RIPK1, ZBP1 and RIPK3 in HT29 / Tet-On-ZBP1 cells treated with 0 or 500 ng/ml Dox for 16h. After enrichment, beads were incubated with 0 or 1 µM USP21 at 30°C for 1 h for deubiquitination. Asterisk indicates unspecific bands. (C) Enrichment of Ub-conjugates for analysis of the ubiquitination status of RIPK1 and ZBP1 using GST-1xUBA or linkage-specific SUBs in HT29 / Tet-On-ZBP1 cells treated with 0 or 500 ng/ml Dox for 16 h. (D) Normalized NF-κB activity in HEK293FT / RIPK3-2xFV cells transfected with GFP (Ctrl) or GFP-tagged SUBs together with EV or ZBP1-expressing plasmids as indicated, co-expressing dual luciferase plasmids. Luciferase activity was measured 24 h after transfection and normalized to GFP (Ctrl)+EV-transfected condition. Data is plotted as mean with S.E.M. (n = 4). One-way ANOVA and Sidak’s multiple comparisons tests were used to test for statistical differences between the indicated condition and ZBP1/GFP-transfected condition. ****, p < 0.0001. Expression levels of ZBP1 and SUBs were analyzed by Western blotting. (E) Relative *CXCL8* mRNA levels in HCT116 / Tet-On-GFP-K63-SUB / RIPK3-2xFV, HCT116 / Tet-On-GFP-M1-SUB / RIPK3-2xFV and HCT116 / Tet-On-GFP / RIPK3-2xFV cells treated with 0 or 100 ng/ml Dox for 48 h before stimulated with 0 or 100 nM dimerizer for 3 h. Data is plotted as mean with S.E.M. (n = 4). Brown-Forsythe and Welch ANOVA tests and Dunnet’s T3 multiple comparisons test were used to test for statistical differences between indicated conditions. ns, p = 0.9997, **, p = 0.0030, *, p = 0.0366. Cells were analyzed by western blotting for the inducible-expression levels. (F, G) Normalized NF-κB activity in HEK293FT / RIPK3-2xFV cells co-expressing ZBP1 with variants of CYLD or OTULIN, as determined by co-transfected dual luciferase reporters. NF-κB activity was measured 24 h after transfection and normalized to ZBP1/EV-transfected cells. Data is plotted as mean with S.E.M. Multiple Welch t-tests were used to test for statistical differences between the indicated condition and the ZBP1/EV-transfected condition. (F) n = 3. *, p = 0.0137, ns, p = 0.8374. (G) n = 4. **, p = 0.0012, ns, p = 0.2342. Cell lysates were loaded for western blot analysis.

Binding of SUBs to the corresponding Ub chains can block their signaling capability in cells (Hrdinka *et al*, 2016; van Wijk *et al*, 2012; Sims *et al*, 2012; Fiil *et al*, 2013). Consistent with our biochemical data, expression of K63-SUB and M1-SUB efficiently inhibited ZBP1-induced NF-κB activity in HEK293FT / RIPK3-2xFV cells and dimerizer-induced expression of *CXCL8* in HCT116/RIPK3-2xFV (Figures 3D and 3E). Also, transient expression of OTULIN, which cleaves M1-Ub, or CYLD, which preferentially cleaves K63- and M1-Ub, inhibited ZBP1-induced NF-κB activity in HEK293FT / RIPK3-2xFV cells (Figures 3F and 3G). Collectively, this shows that K63-Ub and M1-Ub both contribute to ZBP1-RIPK3-dependent inflammatory signaling.

The involvement of M1-Ub implied that LUBAC contributes to ZBP1-RIPK3-dependent inflammatory signaling. Indeed, RIPK3 oligomerization-induced signaling was substantially reduced in HCT116 cells deficient for HOIP (Hrdinka *et al*, 2016), the catalytic subunit of LUBAC, as compared to WT cells (Figure 4A). Also, ZBP1-induced NF-κB activity was substantially impaired by siRNA-mediated knockdown of HOIP in HEK293FT cells (Figure 4B). K63-Ub assembled by cIAPs facilitates the recruitment of LUBAC to the TNF receptor signaling complex (Haas *et al*, 2009), which prompted us to investigate their involvement in ZBP1-RIPK3 inflammatory signaling. Depletion of cIAPs by CpA completely inhibited ZBP1 and RIPK3-oligomerization-induced inflammatory signaling and chemokine production (Figures 4C-4F, S4B and S4C). In addition to depletion of cIAPs, CpA antagonizes the function of XIAP in NOD2 signaling when used at 1 µM or above (Damgaard *et al*, 2013). However, RIPK3 oligomerization-induced inflammatory signaling was comparable in WT and XIAP-knockout HCT116 cells (Figure S4D), indicating that the inhibition of ZBP1 and RIPK3 inflammatory signaling by CpA reflects a requirement for cIAPs.

**Figure 4.**
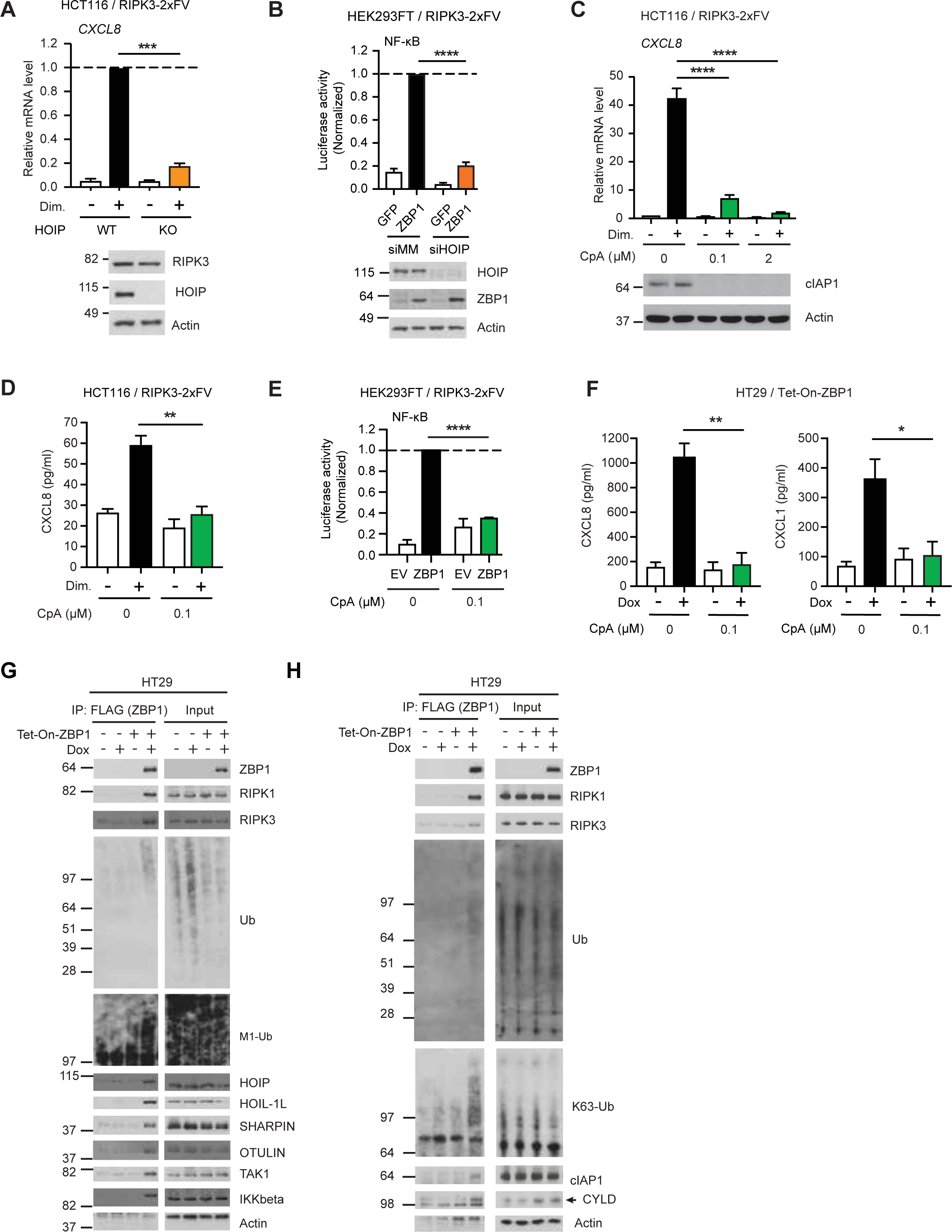
**See also Figure S4. cIAP1 and LUBAC control ZBP1-induced inflammatory signaling** (A) Relative *CXCL8* mRNA levels in WT or HOIP-knockout HCT116 cells stably expressing RIPK3-2xFV and treated with 0 or 100 nM dimerizer for 3 h. Data is plotted as mean with S.E.M. (n = 3). A Welch’s t-test was used to test for the statistical difference as indicated. ***, p = 0.0010. Cell lysates were loaded for western blot analysis. (B) Normalized NF-κB activity in control or HOIP-knockdown HEK293FT / RIPK3-2xFV cells expressing ZBP1. Cells were transfected with siRNA targeting mismatch sequence (siMM) or HOIP. 48 h after siRNA transfection, cells were transfected again with dual luciferase reporters and GFP or ZBP1-expressing plasmids. Luciferase activity in the cell lysate was measured 24 h after the second transfection and normalized to siMM+ZBP1–transfected condition. Data is plotted as mean with S.E.M. (n = 4). A Welch’s t-test was used to test for the statistical difference between indicated conditions. ****, p < 0.0001. HOIP knockdown and ZBP1 expression levels were analyzed by Western blotting. (C) Relative *CXCL8* mRNA levels in HCT116 / RIPK3-2xFV cells pretreated with CpA for 0.5 h before treated with 0 or 100 nM dimerizer for 3 h. Data is plotted as mean with S.E.M. (n = 3). One-way ANOVA and Sidak’s multiple comparisons test were used to test for statistical differences between the indicated condition and the dimerizer-treated condition. ****, p < 0.0001. Cell lysates were analyzed by western blotting for cIAP1 levels. (D) CXCL8 concentration in the cell culture supernatant of HCT116 / RIPK3-2xFV cells pretreated with 100 nM CpA for 1 h before stimulated with 0 or 100 nM dimerizer for 24 h. Data is plotted as mean with S.E.M. (n = 3). An unpaired t-test was used to test for the statistical difference between indicated conditions. **, p = 0.0042. (E) Normalized NF-κB activity in HEK293FT / RIPK3-2xFV cells treated with 0 or 100 nM CpA and transfected with EV or ZBP1-expressing vector, co-transfecting dual luciferase reporter plasmids. Data is plotted as mean with S.E.M. (n =3). A Welch’s t-test was used to test for the statistical difference between indicated conditions. ****, p < 0.0001. (F) CXCL8 and CXCL1 levels in cell culture supernatant of HT29 / Tet-On-ZBP1 cells treated with 0 or 100 nM CpA and 0 or 50 ng/ml Dox as indicated for 24 h. Data is plotted as mean with S.E.M. (n = 3). Unpaired t-tests were used to test for the statistical difference between indicated conditions. **, p = 0.0049, *, p = 0.0394. (G, H) Western blot analyses of anti-FLAG (ZBP1) immunoprecipitation from HT29 or HT29 / Tet-On-ZBP1 cells treated with 0 or 500 ng/ml Dox for 16 h.

Consistent with our signaling data, analysis of the ZBP1 complex immunoprecipitated from Dox-treated HT29 cells showed that ZBP1, in addition to RIPK1 and RIPK3, co-immunoprecipitated K63- and M1-Ub, cIAP1, LUBAC (HOIP, HOIL-1, SHARPIN), the LUBAC-associated DUBs OTULIN and CYLD, TAK1, and IKKβ (Figures 4G and 4H). This suggests that ZBP1 forms a pro-inflammatory receptor signaling complex that, akin to other immune receptor complexes, consists of adaptor kinases, K63- and M1-Ub ligases and DUBs, and the ubiquitin-dependent kinases TAK1 and IKKβ.

### RIPK1 is required for ZBP1 signaling

Prompted by our data showing ZBP1-induced RIPK1 ubiquitination, we sought to determine the functional role of RIPK1 in the ZBP1 complex. For this, we generated RIPK1-KO HT29 single cell clones and integrated the Dox-inducible ZBP1 expression system. Strikingly, both ZBP1-induced chemokine production and cell death were blocked in the absence of RIPK1, demonstrating a central role of RIPK1 in mediating signaling responses by ZBP1 in HT29 cells (Figures 5A and 5B, S5A and S5B).

**Figure 5.**
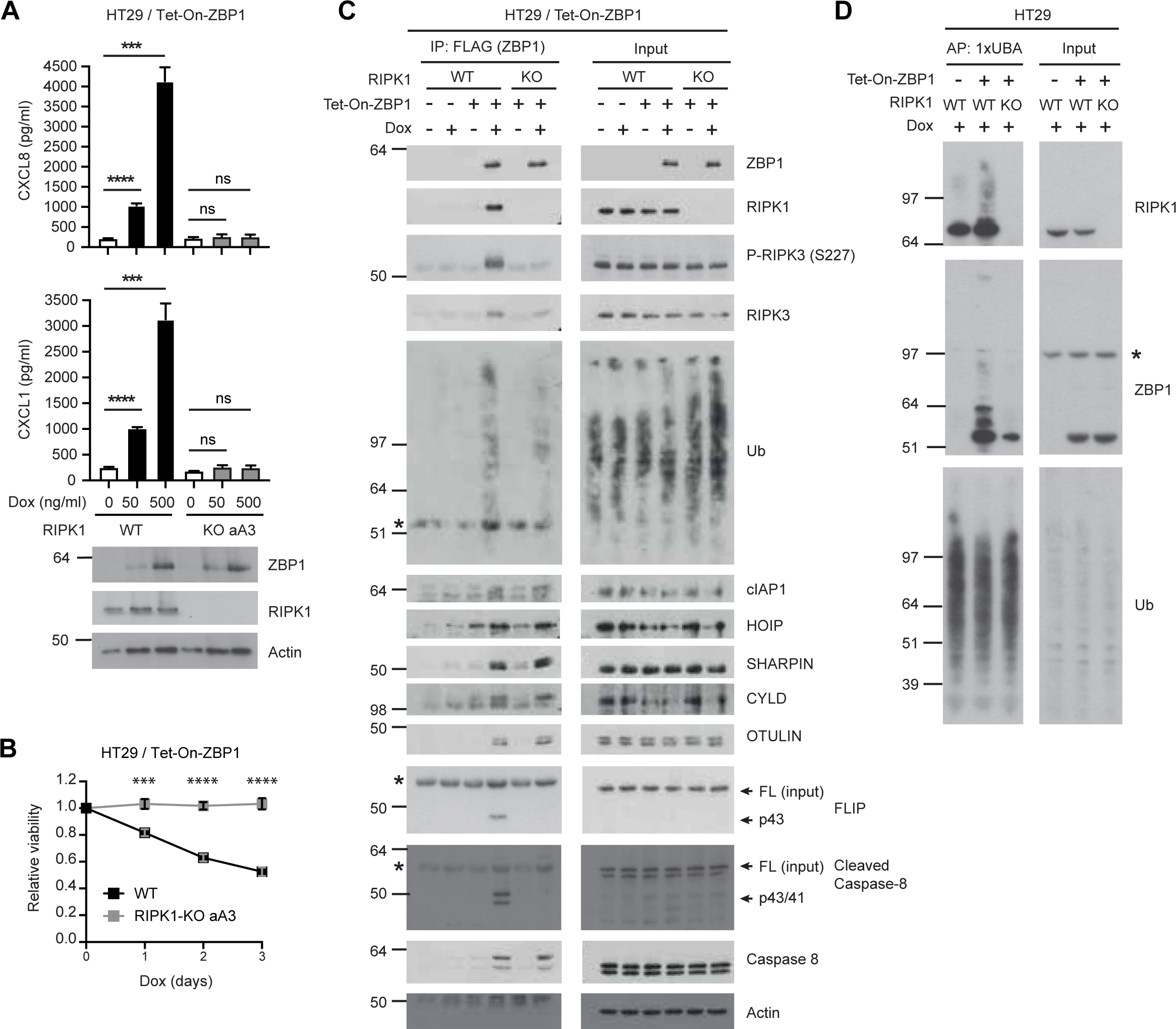
**See also Figure S5. RIPK1 is required for ZBP1 signaling** (A) ELISA analyses of chemokine concentrations in the supernatants of HT29 / Tet-On-ZBP1 or HT29 / RIPK1-KO clone aA3 / Tet-On-ZBP1 cells treated with 0, 50 or 500 ng/ml Dox. data is plotted as mean with S.E.M. (n = 6). Brown-Forsythe and Welch ANOVA tests and Dunnet’s T3 multiple comparisons test were used to test for statistical differences between indicated conditions. Cell lysates were analyzed by western blotting for ZBP1 levels. ns, p > 0.2, ***, p = 0.0003 for CXCL8, p = 0.0008 for CXCL1, ****, p < 0.0001. (B) Relative viability of HT29 / Tet-On-ZBP1 or HT29 / RIPK1-KO clone aA3 / Tet-On-ZBP1 cells treated with 500 ng/ml Dox for 3, 2, 1 or 0 day(s). Measurement values are normalized to 0 day of treatment within each cell line. Data is plotted as mean with S.E.M. (n = 3). Two-way ANOVA and Sidak’s multiple comparison tests were used to test for statistical differences between WT and RIPK1-KO cells at the indicated time points after Dox treatment. ***, p = 0.0003, ****, p < 0.0001. (C) Western blot analyses of anti-FLAG (ZBP1) immunoprecipitation from HT29, HT29 / Tet-On-ZBP1 or HT29 / RIPK1-KO clone aA3 / Tet-On-ZBP1 cells treated with 0 or 500 ng/ml Dox for 16 h. Asterisk indicates antibody heavy chain signal in PD samples. (D) TUBE pull down analyses of the ubiquitination status of RIPK1 and ZBP1 from HT29, HT29 / Tet-On-ZBP1 or HT29 / RIPK1-KO aA3 / Tet-On-ZBP1 cells treated with 500 ng/ml Dox for 16 h. Asterisk indicates antibody background signal.

The association of RIPK3 with ZBP1, especially its phosphorylated form, was greatly reduced in the absence of RIPK1, indicating a primary function of RIPK1 in stabilizing the ZBP1-RIPK3 interaction (Figure 5C). Intriguingly, the association of ZBP1 with LUBAC, cIAPs, CYLD and OTULIN was not affected in RIPK1-KO cells as compared to WT cells, while there was a reduction in co-immunoprecipitated Ub-conjugates with ZBP1 (Figure 5C). Correspondingly, the ubiquitination on ZBP1 was substantially reduced in RIPK1-KO cells (Figure 5D). This supported the notion that the ubiquitination of RIPK1 and ZBP1 is important for ZBP1-dependent inflammatory signaling, although it is not clear how RIPK1 regulates ZBP1 ubiquitination.

### RIPK1 links caspase-8 to RIPK3 to suppress kinase activity-dependent inflammatory signaling

Strikingly, p43/41-caspase-8 and p43-cFLIP_L_, processed as part of a catalytically active heterodimer, were co-immunoprecipitated with ZBP1 in a RIPK1-dependent manner (Figure 5C). Since RIPK1 links caspase-8 to RIPK3 to suppress necroptosis, we speculated that caspase-8 might regulate ZBP1-RIPK3 inflammatory signaling. Indeed, zVAD treatment enhanced the production of CXCL8 by Dox-induced ZBP1 expression in HT29 cells and chemokine expression following dimerizer treatment of HCT116/RIPK3-2xFV cells (Figures 6A and 6B). Also, shRNA-mediated knockdown of caspase-8 in HCT116/RIPK3-2xFV cells led to increased dimerizer-induced chemokine expression (Figure 6C). Strikingly, the enhanced signaling in the absence of caspase-8 activity was inhibited by GSK’872, Nec1s and NSA, as well as by TAK1 and IKK inhibitors (Figures 6A-6C and S6A). These data show that in the absence of caspase-8 activity, ZBP1-induced inflammatory signaling is, in part, mediated by RIPK3 and RIPK1 kinase activity as well as MLKL.

**Figure 6.**
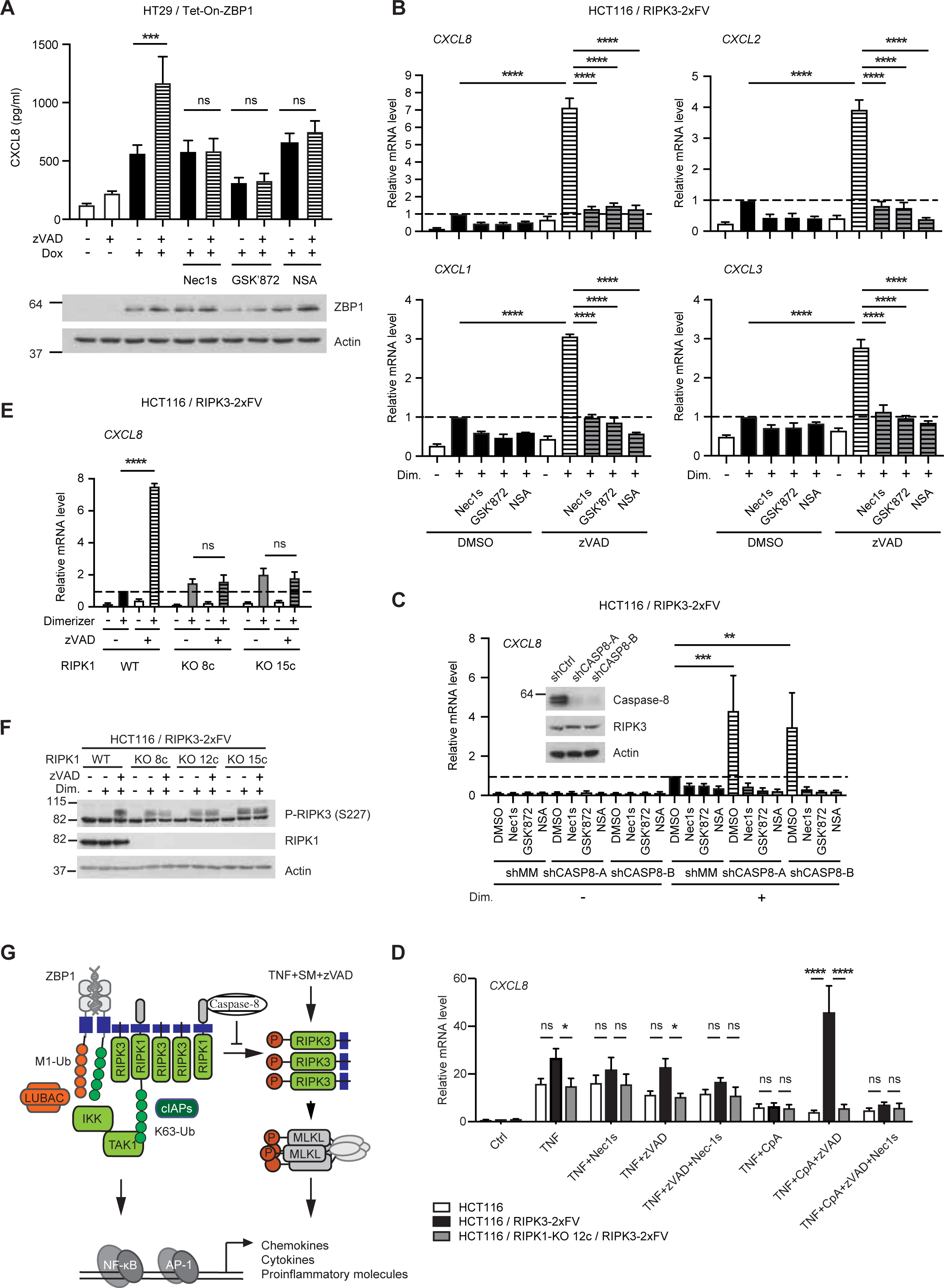
**See also Figure S6. RIPK1 links caspase-8 to RIPK3 to suppress kinase activity-dependent inflammatory signaling** (A) CXCL8 concentrations in the supernatants of HT29 / Tet-On-ZBP1 cells treated with 0 or 500 ng/ml Dox in combination with DMSO, 10 µM Nec1s, 10 µM GSK’872 or 1 µM NSA for 24 h. Data is plotted as mean with S.E.M. (n = 4). One-way ANOVA and Sidak’s multiple comparisons tests were used to test for statistical differences between indicated conditions. ***, p = 0.0005, ns, p > 0.9999 for Nec1s, p = 0.9995 for GSK’872, p = 0.7950 for NSA. Cell lysates from the same experiment were analyzed by western blot for the expression level of ZBP1. (B) Relative chemokine mRNA levels in HCT116 / RIPK3-2xFV cells pretreated with 0 or 20 µM zVAD in combination with DMSO, 10 µM Nec1s, 10 µM GSK’872 or 1 µM NSA for 1 h, followed by treatment with 0 or 100 nM dimerizer for 3 h. Data is plotted as mean with S.E.M. (n = 3). ****, p < 0.0001. One-way ANOVA and Sidak’s multiple comparison test were used to test for statistical differences between indicated conditions. (C) Relative *CXCL8* mRNA levels in HCT116 / RIPK3-2xFV cells stably knocked down against mismatch (shMM) or two different sites of caspase-8 (shCASP8-A and shCASP8-B), pretreated with DMSO, 10 µM Nec1s, 10 µM GSK’872 or 1 µM NSA for 1h followed by treatment with 0 or 100 nM dimerizer for 3h. Data is plotted as mean with S.E.M. (n = 3). Two-way ANOVA and Tukey’s multiple comparison tests were used to test for statistical differences between indicated conditions. ***, p = 0.0001, **, p = 0.0035. Equal number of cells was collected from each cell line for western blot analysis of caspase-8 knockdown levels. (D) Relative *CXCL8* mRNA levels in HCT116, HCT116 / RIPK3-2xFV or HCT116 / RIPK1-KO clone 12c/RIPK3-2xFV cells pretreated with combinations of 2 µM CpA, 20 μM zVAD and 10 μM Nec1s as indicated for 1h, and then stimulated with 0 or 2 ng/ml TNF for 3 h. Data is plotted as mean with S.E.M. (n = 3). Two-way ANOVA and Tukey’s multiple comparison tests were used to test for statistical differences between indicated conditions. ns, p > 0.05, *, p = 0.0419 in TNF-treated condition, p = 0.0317 for TNF+zVAD–treated condition, ****, p < 0.0001. (E) Relative *CXCL8* mRNA levels in WT or RIPK1-KO clones of HCT116 cells stably expressing RIPK3-2xFV, pretreated with 0 or 20 µM zVAD for 1 h before treated with 0 or 100 nM dimerizer for 3 h. Data is presented as mean with S.E.M. (n = 4). One-way ANOVA and Sidak’s multiple comparisons test were used to test for statistical differences between indicated conditions. ****, p < 0.0001, ns, p = 0.9836 for clone 8c, p = 0.8750 for clone 15c. (F) Western blot analyses of RIPK3 in WT and three independent RIPK1-knockout clones of HCT116 cells stably expressing RIPK3-2xFV, pretreated with 0 or 20 μM zVAD for 1 h before treated with 0 or 100 nM dimerizer for 6 h. (G) Schematic illustration of RIPK3-mediated inflammatory signaling pathways.

This was reminiscent of previous reports showing that necroptotic signaling stimulated by TNF in the presence of Smac mimetics and caspase inhibitor zVAD (TSZ) induces RIPK1-RIPK3-MLKL-dependent expression of chemokines concomitantly with cell death (Zhu *et al*, 2018). In concordance, TSZ (TNF+CpA+zVAD) stimulated strong *CXCL8* expression in HCT116 / RIPK3-2xFV cells but not in parental HCT116 cells, whereas TNF-, TNF+CpA-, and TNF+zVAD-induced *CXCL8* expression was largely independent of RIPK3 (Figure 6D). No cell death was detected following TSZ treatment at the time of the assay, demonstrating that the RIPK3-inflammatory signaling pathway is engaged by TNF without cell death (Figure S6B). Akin to ZBP1 signalling when caspase-8 is inhibited, TSZ-induced *CXCL8* expression in HCT116 / RIPK3-2xFV cells was inhibited by Nec1s, RIPK1-KO, GSK’872, kinase-dead RIPK3 (K50A or D142N), NSA or knockdown of MLKL (Figures 6D, S6C and S6D).

To determine if RIPK1 is required for caspase-8 to suppress RIPK3 kinase activity-mediated signaling, we treated HCT116 / RIPK1-KO cells with zVAD while inducing RIPK3 oligomerization. Whilst RIPK1 was not needed for RIPK3 oligomerization-induced chemokine expression, RIPK1 was required for the enhancement of chemokine expression by zVAD (Figure 6E). Under caspase-inhibited conditions, RIPK3 oligomerization led to the accumulation of phosphorylated RIPK3 (P-RIPK3) that migrated slower on SDS-PAGE than the unmodified RIPK3 (Figure 6F). This slower-migrating P-RIPK3 likely represents the kinase-activated form of RIPK3, which, in the absence of zVAD, is suppressed by caspase-8 (He *et al*, 2009; Choi *et al*, 2018). This is supported by the appearance of a ∼30 kDa RIPK3 fragment after dimerizer-treatment of cells expressing RIPK3-2xFV (Figures 1F and S2H; (Feng *et al*, 2007)). Consistent with the idea that RIPK1 links caspase-8 to RIPK3, dimerizer treatment of RIPK1-KO cells induced the accumulation of the slower-migrating P-RIPK3 form even without caspase inhibition (Figure 6F).

Together, these data show that caspase-8 activity restricts ZBP1 inflammatory signaling by inhibiting RIPK3 kinase activity-mediated signaling and suggests that RIPK1 mediates the suppression of kinase-activated RIPK3 (Figure 6G).

### ZBP1 mediates SARS-CoV-2-induced cytokine production

Lastly, we investigated the contribution of ZBP1-induced inflammatory responses during virus infection. COVID-19, caused by the SARS-CoV-2 virus, is characterized by lower antiviral responses and high inflammatory chemokine and cytokine production (Blanco-Melo *et al*, 2020). Analysis of publicly available RNA-sequencing datasets (Blanco-Melo *et al*, 2020) showed a substantial increase in *ZBP1* expression in post-mortem patient lung biopsies compared to healthy controls (Figure 7A). In addition, analysis of a single-cell RNA-sequencing dataset (Ren *et al*, 2021) showed that ZBP1 expression was significantly higher in COVID-19 patients in the progressive disease stage than those in the convalescent stage or healthy controls (Figure 7B). Monocytes and neutrophils have been found to be two major sources of cytokines in the peripheral blood that contribute to the “cytokine storm” in critically ill COVID-19 patients (Ren *et al*, 2021; Vanderbeke *et al*, 2021). Analysis of the single-cell RNAseq dataset (Ren *et al*, 2021) revealed an increase in ZBP1 expression in neutrophils and all major inflammatory subsets of monocytes in COVID-19 patients in progressive stage compared to convalescent stage patients (Figure S7A). Taken together, these analyses indicate a potential role of ZBP1 in fueling the cytokine storm in COVID-19 patients.

**Figure 7.**
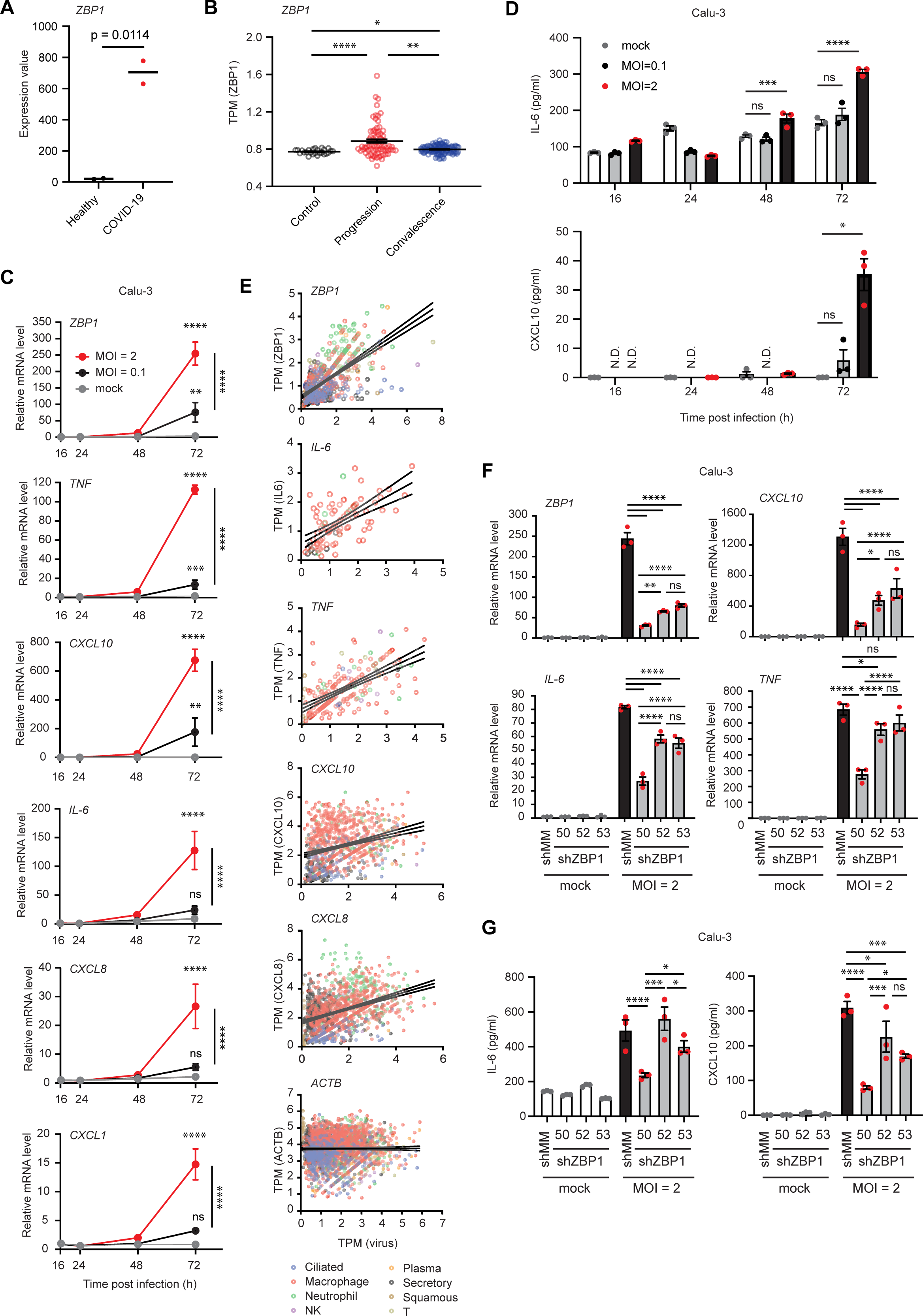
**ZBP1 mediates SARS-CoV-2-induced inflammation** (A) Expression values of *ZBP1* in post-mortem lung samples of two COVID-19 patients compared with healthy lung biopsies (Blanco-Melo *et al*, 2020). An unpaired t-test was used to test for statistical differences between the two conditions. (B) Patient averaged single cell Transcript Per Million (TPM) values of *ZBP1* in lung and peripheral blood of COVID-19 patients in progressive or convalescent stage, compared to healthy controls (Ren *et al*, 2021). Data is plotted as individual values per patient with mean and S.E.M. (n = 25 for healthy, n = 77 for progression, n = 102 for convalescence patients). Kruskai-Wallis test and Dunn’s multiple comparisons test were used to test for statistical differences between indicated conditions. ****, p < 0.0001, *, p = 0.0375, **, p = 0.0036. (C) Relative mRNA levels of indicated genes in Calu-3 cells infected with mock or SARS-CoV-2 viruses at indicated MOIs for 16, 24, 48 or 72 hours. Data is plotted as mean with S.E.M. (n = 3). Two-way ANOVA and Tukey’s multiple comparison test were used to test for statistical differences between each infected sample and mock-treated sample and between the two different MOIs at 72 h time point. ****, p < 0.0001, ***, p = 0.0006, **, p = 0.0022 for *ZBP1*, p = 0.0072 for *CXCL10*, ns, p = 0.6080 for *IL*-6, p = 0.5842 for *CXCL8*, p = 0.1243 for *CXCL1*, (D) Cytokine levels in cell culture supernatant of Calu-3 cells infected with mock or SARS-CoV-2 viruses at indicated MOIs for 16, 24, 48 or 72 hours. Data is plotted as mean with S.E.M. (n = 3). Two-way ANOVA and Dunnet’s multiple comparisons test were used to test for statistical differences between indicated conditions. ns, p = 0.6762 for IL-6 at 48 h, p = 0.0722 for IL-6 at 72 h, p = 0.6378 for CXCL10 at 72 h, ****, p < 0.0001, ***, p = 0.0002, *, p = 0.0244. (E) Pearson’s correlation of individual single cell TPM values of indicated genes with virus load (Ren *et al*, 2021). Data is presented as individual values from each cell, with linear regression line and its 95% confidence bands. *ZBP1*, n = 720, *IL*-6, n = 94, *TNF*, n = 211, *CXCL10*, n = 1053, *CXCL8*, n = 1463, *ACTB*, n = 2683 (cells). The positive correlation is significant (p < 0.0001) between TPM (virus) and TPM (*ZBP1*), TPM (*IL*- 6), TPM (*TNF*), TPM (*CXCL10*) or TPM (*CXCL8*). Correlation between TPM (virus) and TPM (*ACTB*) is not significant (p = 0.7862). (F, G) Relative mRNA levels (F) or secreted cytokine levels (G) of indicated genes in Calu- 3 cells knocked down against mismatch sequence (shMM) and ZBP1 at three different sites (shZBP1-50, shZBP1-52 and shZBP1-53) infected with mock or SARS-CoV-2 virus at MOI = 2 for 72 hours. Data is presented as mean with S.E.M. (n = 3). One-way ANOVA and Sidak’s multiple comparisons test were used to test for statistical differences between indicated conditions. ns, not significant, *, p<0.05, **, p<0.01, ***, p<0.001, ****, p<0.0001. See source data for statistical details.

To investigate the role of ZBP1 in SARS-CoV-2 infection, Calu-3 human lung adenocarcinoma cells were infected at two different multiplicities of infection (MOI), and the expression profiles of a panel of inflammatory genes were determined over time. This showed that SARS-CoV-2, in a dose-dependent manner, caused the upregulation of *ZBP1*, cytokines *IL-6* and *TNF*, and chemokines *CXCL10*, *CXCL8* and *CXCL1* between 48 h and 72 h after infection, and led to the secretion of IL-6 and CXCL10 (Figures 7C and 7D). In concordance, analysis of the single-cell RNA sequencing dataset revealed positive correlations between virus load and the expression levels of ZBP1 and various cytokines in virus-positive cells from bronchoalveolar lavage fluid (BALF) and sputum samples (Figure 7E).

We then established three independent Calu-3 cell lines with stable knockdown of *ZBP1* (shZBP1) and control cells expressing a non-targeting shRNA (shMM; Figure 7F, top-left panel). Infection with SARS-CoV-2 (MOI=2) showed that knockdown of *ZBP1* attenuated the expression of the measured cytokine and chemokine genes, as well as the production of CXCL10 and to a lesser degree of IL-6 (Figures 7F and 7G). The attenuation of the pro-inflammatory response after infection correlated with the efficacy of the individual shRNAs to silence *ZBP1* expression; shZBP1-50 resulted in a substantially stronger attenuation than did shZBP1-52 or shZBP1-53 (Figures 7F and 7G). The effect of ZBP1 knockdown was less prominent for interferon-induced genes (Figure S7B), and the intracellular virus amount 3 days after infection was similar in all cell lines (Figure S7C). Of note, whilst ZBP1 expression can trigger cell death, the corresponding markers remained negative by Western blotting analysis in Calu-3 cells up to 3 days after SARS-CoV-2 infection (Figure S7D). Together, this suggests that ZBP1, subsequent to its transcriptional upregulation by SARS-CoV-2 infection, stimulates the production of inflammatory chemokines and cytokines that contribute to the severity of COVID-19 symptoms.

Altogether, our data reveal that ZBP1 can activate Lys63- and Met1-Ub-dependent inflammatory signalling mediated by a kinase-independent scaffolding function of RIPK3 and RIPK1, and suggest that this pathway contributes to inflammatory responses triggered by SARS-CoV-2.

## Discussion

### RIPK3 as a scaffolding kinase for inflammatory signaling

RIPK3 was initially reported to stimulate both inflammatory signaling and cell death, but RIPK3-deficient cells showed normal NF-κB signaling in response to the stimulation of TNFR1, B and T cell receptors, and Toll-like receptors (TLRs) 2 and 4, excluding the role of RIPK3- mediated inflammatory signaling in those contexts (Yu *et al*, 1999; Newton *et al*, 2004; Sun *et al*, 1999). Here, we demonstrate that RIPK3 is a *bona fide* inflammatory mediator by identifying it as an essential adaptor protein in ZBP1-dependent inflammatory signaling in human cells. Interestingly, the role of RIPK3 in inflammatory signaling depends on its RHIM domain and does not require its kinase activity, which is in contrast to its role in necroptosis where kinase activity is essential. This indicates that RIPK3 acts as a scaffold and not as a kinase in ZBP1- induced inflammatory signaling.

This is reminiscent of the scaffolding role of other receptor-associated kinases in mediating inflammatory signaling, including RIPK1 in TNFR1 signaling, RIPK2 in NOD2 signaling, and IL-1R-associated kinases (IRAKs) in IL-1R signaling (Ea *et al*, 2006; Hrdinka *et al*, 2018; Ordureau *et al*, 2008; Koziczak-Holbro *et al*, 2008). While these kinases serve as scaffolds for the formation of K63- and/or M1-Ub, ubiquitination of RIPK3 was not consistently detected in response to ZBP1 expression. Instead, ZBP1 and RIPK1 were both modified with K63- and/or M1-Ub. Together with the requirement of RIPK1 and the RHIM of RIPK3 for ZBP1 inflammatory signaling, this suggests that RIPK3 may mediate signaling by RHIM-mediated recruitment and/or stabilization of RIPK1 in the ZBP1 complex, whereas RIPK1 and ZBP1 are the primary ubiquitination targets to facilitate NF-κB activation. Since the deletion of RIPK1 reduced the association of RIPK3, in particular its phosphorylated form, with ZBP1, RIPK1 may also contribute to the stabilization of a RHIM-mediated ZBP1-RIPK3-RIPK1 complex. This would be concordant with previous reports showing that a RIPK1-RIPK3 interaction precedes formation of RIPK3 oligomers during the activation of RIPK3 (Li *et al*, 2012; Wu *et al*, 2014). Of note, our study of the ZBP1 signaling complex relied on Dox-induced overexpression of ZBP1, which precludes a detailed time-resolved study of the assembly of the signaling complex and of ubiquitination dynamics of complex components in response to ligand-binding. Such investigations will be warranted when cognate ZBP1 ligands are better defined in order to gain detailed insights into the assembly of the ZBP1 signaling complex.

It was intriguing that RIPK3, in addition to its scaffolding role, also promoted inflammatory signaling by ZBP1 in a kinase activity-dependent manner when caspase-8 activity was inhibited. This is reminiscent of the inflammatory signaling pathway described for TNF-induced necroptosis (TSZ treatment) in HT29 cells, which is mediated by RIPK3 and RIPK1 kinase activity and by MLKL (Zhu *et al*, 2018). This suggests that caspase-8 represents a checkpoint switch for RIPK3 kinase activity-mediated signaling also in the context of ZBP1 by constantly suppressing the kinase activity-dependent inflammatory signaling pathway after the engagement of RIPK3. While it remains to be defined how the RIPK3 kinase activity-dependent signaling leads to inflammatory gene activation, our observations indicate that the default role of RIPK3 in ZBP1 inflammatory signaling is as a scaffolding kinase and that the kinase activity-dependent pathway is activated when caspase-8 activity is antagonized, such as during infection by viruses encoding caspase inhibitors.

It is not fully understood how caspase-8 regulates RIPK3 kinase activity-dependent signaling but our data suggest that caspase-8 activity suppresses kinase-activated RIPK3. This notion is supported by the RIPK1-dependent co-immunoprecipitation of both the p43/41-caspase-8- p43-FLIP heterodimer and phosphorylated RIPK3 with ZBP1, and is in concordance with previous findings that kinase-activated RIPK3 is recognized and depleted from the cytosolic pool (Choi *et al*, 2018).

### ZBP1-mediated inflammatory signaling and cell death responses

Since the discovery that ZBP1, via RIPK3, induces necroptosis during murine cytomegalovirus (MCMV) infection, its role in cell death during infection and embryonic development has been well established (Kuriakose & Kanneganti, 2018; Upton *et al*, 2012; Lin *et al*, 2016; Thapa *et al*, 2016; Wang *et al*, 2020; Newton *et al*, 2016; Jiao *et al*, 2020). Our study expands the understanding of ZBP1’s function as we uncover that ZBP1, in human cell lines, contributes to SARS-CoV-2-induced chemokine and cytokine expression, and that ZBP1, in a ligand binding-dependent manner, triggered RIPK3-mediated inflammatory signaling at a lower expression threshold and at earlier time points than needed for stimulation of cell death. It is tempting to speculate that ZBP1, akin to other innate immune receptors, induces ubiquitin-dependent NF-κB signaling as a first line of defense to recruit innate immune cells (e.g. neutrophils and monocytes), and that triggering of cell suicide is a backup mechanism invoked during pathological conditions where ZBP1 expression is highly induced by interferons. Curiously, the major mode of cell death in ZBP1-expressing HT29 cells appeared to be RIPK1- mediated apoptosis whereas previous studies in murine systems showed that RIPK1 restricts ZBP1-RIPK3-induced necroptosis during development and in skin inflammation (Lin *et al*, 2016; Newton *et al*, 2016; Devos *et al*, 2020). Whether this represents a difference between mice and human or is specific to the experimental systems is interesting and should be addressed in future studies.

### ZBP1 and SARS-CoV-2

Our SARS-CoV-2 experimental data and analysis of RNAseq datasets from COVID-19 patients show that ZBP1 is upregulated by SARS-CoV-2 infection and that its expression correlates with the expression of pro-inflammatory chemokines and cytokines, including CXCL10/IP-10 and IL-6 (Figure 7). IL-6 and CXCL10 are upregulated in critically ill COVID-19 patients as part of the cytokine storm, and IL-6 levels correlate with mortality (Ruan *et al*, 2020; Huang *et al*, 2020; Chen *et al*, 2020). This suggests that ZBP1-mediated inflammatory signaling may contribute to an unbalanced immune response in COVID-19 patients, characterized by reduced antiviral signaling and enhanced inflammatory levels (Blanco-Melo *et al*, 2020).

SARS-CoV-2-induced ZBP1 expression in Calu-3 cells did not lead to detectable cell death, supporting the notion that ZBP1, as a primary response, stimulates inflammatory signaling in human cells. However, our observation does not exclude the possibility of ZBP1-induced cell death during SARS-CoV-2 infections *in vivo*, where ZBP1 expression may be further induced by interferons produced by immune cells (Fu *et al*, 1999; Takaoka *et al*, 2007). Conversely, it is plausible that ZBP1-induced inflammatory signaling contributes to pathological inflammatory conditions where ZBP1-mediated cell death has been reported (Upton *et al*, 2012; Kuriakose *et al*, 2016; Thapa *et al*, 2016; Newton *et al*, 2016; Lin *et al*, 2016; Jiao *et al*, 2020). Irrespective, our data suggest that antagonizing ZBP1 signaling pharmacologically could attenuate pathological inflammation.

## Materials and Methods

### Lead Contact

Further information and requests for resources and reagents should be directed to and will be fulfilled by the Lead Contact Prof. Mads Gyrd-Hansen.

### Materials Availability

Plasmids and cell lines generated in this study are available from lead contact with a completed Materials Transfer Agreement.

### Cell lines

All cell lines were cultured at 37°C and 5% CO_2_ in growth medium supplemented with 10% v/v fetal bovine serum (FBS, Labtech FCS-SA), 60 μg/ml penicillin and 100 μg/ml streptomycin (PS, Thermo Fisher 15070). The growth medium for HT29 and HCT116 cells was McCoy’s 5A (Modified) (Thermo Fisher 26600), for HEK293FT and Phoenix-Ampho was DMEM (Thermo Fisher 31966-021), for THP1 cells was RPMI (Thermo Fisher 42401042) supplemented with GlutaMAX (Thermo Fisher 35050061), Sodium Pyruvate (Gibco 11360-039) and 50 µM 2-mercaptoethanol (Gibco 31350-010), and for Calu-3 cells was MEM (Thermo Fisher 11090081) supplemented with GlutaMAX, Sodium Pyruvate and non-essential amino acids (Thermo Fisher 11140035). Cells were routinely checked for *Mycoplasma* Spp. contamination with the MycoAlert Mycoplasma Detection kit (Lonza LT-07). Cell lines were not authenticated.

### Isolation of primary neutrophils

Primary human neutrophils were obtained from healthy donors with their written informed consent by The Oxford Radcliffe Biobank with project number ORB 20/A136. The study is authorized by South Central – Oxford C Research Ethics Committee (Ref# 19/SC/0173). Neutrophils were isolated from 50 ml Ficoll-layered blood cone using EasySep Human Neutrophil Isolation Kit (StemCell 17957) following manufacturer’s instructions.

### Generation of cell lines stably expressing RIPK3 variants

For the construction of HT29- and HCT116-based RIPK3-expressing cell lines, LZRS-zeo- based retroviral vectors were used (Rodriguez *et al*, 2016). To produce retroviral particles, Phoenix-Ampho cells were plated at a density of 3.5x10^6^ cells in 10 cm dishes in 15 ml complete growth media. The next day, transfection was carried out with a mixture of 1.5 ml OptiMEM, 36 μl FuGENE HD (Promega E2311), 1.2 μg pMD.G (VSVG) plasmid and 10.8 µg of the retroviral vector, and media was replaced 24h after transfection. Two batches of retroviral particles were collected, respectively at 48 and 72 hours post-transfection. Retroviral particles were precipitated by incubating 0.45 µm syringe-filtered culture media in 150 mM NaCl and 5% PEG-8000 at 4°C overnight, followed by centrifugation at 3500 x*g* for 15 min. Pellets were resuspended in 150 µl sterile PBS and stored at -80°C until use.

For retroviral transduction, between 1 and 3x10^5^ cells were seeded into 6-well plates. The next day, cells were transduced using 1 ml virus-containing supernatant or 25 µl precipitated viral particles in the presence of 10 µg/ml polybrene in a total of 2 ml complete growth medium. Cells were incubated with virus-containing supernatant for 24 h before replaced with complete growth medium to rest. 48-72 hours after transduction, cells were selected with 250 ng/µl zeocin (Invitrogen R250) until two passages after the complete elimination of non-transduced cells in the control well.

### Generation of HCT116 cells with inducible GFP-SUB expression

To generate HCT116 / Tet-On-GFP-SUB / RIPK3-2xFV cells, HCT116 cells were first transduced with lentiviral particles generated from pLenti-CMV-Blast plasmids carrying the Tet Repressor gene. During selection with 5 µg/ml blasticidin (Thermo Fisher A1113903), single clones were isolated, and clone C4 with high levels of Tet-Repressor expression was used for the next steps.

HCT116 / Tet-On clone C4 was plated at a density of 5x10^4^ cells per well in 12-well plates, and transduced with 3 µl precipitated lentiviral particles generated with pLVX-tight-puro plasmids encoding GFP, GFP-K63-SUB or GFP-M1-SUB sequences (Hrdinka *et al*, 2016). After selection with 1 µg/ml puromycin (Invitrogen ant-pr) in the presence of 5 µg/ml blasticidin, HCT116 / Tet-On-GFP, HCT116 / Tet-On-GFP-K63-SUB and HCT116 / Tet-On-GFP-M1-SUB cells were transduced with retroviral particles produced from LZRS-zeo-RIPK3-2xFV plasmids and selected with the combination of 5 µg/ml blasticidin, 1 µg/ml puromycin and 250 ng/µl zeocin.

### Production of lentiviral particles

For the production of lentiviral particles, HEK293FT cells were plated at a density of 3.5x10^6^ cells in 10 cm dishes in 15 ml complete growth media. The next day, they were transfected with a mixture of 1.5 ml OptiMEM, 36 μl FuGENE HD, 6 μg psPAX2 vector, 1.5 μg pMD.G (VSVG) and 4.5 μg lentiviral vector. 24 h after transfection, transfection reagent-containing media were replaced with 10 ml complete growth media to allow the secretion of lentiviral particles for 72 h. Virus-containing supernatant was filtered, and lentiviral particles were precipitated as described for retroviral particles or directly frozen at -80°C for preservation.

### Generation of HT29 cells with Dox-inducible expression of ZBP1

For lentivirus production, HEK293T cells were transfected with C-terminally FLAG-tagged wild-type or Zα1α2-mutant human ZBP1-expressing transducing vectors in the doxycycline-inducible Tet-On pDG2 backbone (De Groote *et al*, 2016) together with the pCMV delta R8.91 gag-pol–expressing packaging plasmids and pMD2.G VSV-G-expressing envelope plasmid. HT29 and HT29 / RIPK1-KO clones were transduced using 100 µl 0.45 µm syringe-filtered lentivirus-containing supernatant in 12-well plate. 72 h after transduction, cells were selected with puromycin as described above.

### Generation of shRNA-mediated stable knockdown cell lines using lentiviral particles

Lentiviral particles were used to construct stable cell lines knocked down against mismatch control sequence (shMM, Sigma SHC002) or target genes. shRNA plasmids used for this study are pLKO.1-based targeting the following sequences:

shCASP8-A (Sigma SHCLND, TRCN0000377309): CACCAGGCAGGGCTCAAATTT; shCASP8-B (Sigma SHCLND, TRCN0000376481): GGAGCTGCTCTTCCGAATTAA; shZBP1-50 (Sigma SHCLNG TRCN0000123050): GCACAATCCAATCAACATGAT; shZBP1-52 (Sigma SHCLNG TRCN0000123052): CCACATGAAATCGTGCTTTCT; shZBP1-53 (Sigma SHCLNG TRCN0000123053): CCAAGTCCTCTACCGAATGAA.

For the transduction of lentiviral particles, HCT116 / RIPK3-2xFV cells were seeded in 12-well plates at a density of 0.5-1x10^5^ cells per well and transduced by incubation with 10-20 μl precipitated virus in 1 ml complete growth medium containing 10 μg/ml polybrene for 24 h. Calu-3 cells were seeded in 10 cm dishes at a density of 7.5x10^5^ cells per dish, and transduced by incubation with 750 µl virus-containing supernatant in 10 ml complete growth medium containing 4 μg/ml polybrene for 24 h. 72 h after transduction, cells were selected with puromycin as described above.

### Construction of RIPK1 knockout cell lines by CRISPR/Cas9

Three different RIPK1 CRISPR/Cas9 knockout plasmids (Santa Cruz sc-400377) respectively encoding guide RNA targeting sequences GGCTTTGCGTTGACGTCATTC (gRNAa), GCTCGGGCGCCATGTAGTAG (gRNAb) and CGGCTTTCAGCACGTGCATC (gRNAc) were transfected separately into HCT116 cells in a mixture of 200 µl OptiMEM, 8 µl FuGENE 6 with 2 µg plasmids, or into HT29 cells using a mixture of 500 µl OptiMEM, 20 µl Lipofectamine LTX (Invitrogen 15338100), 3.75 µg of each plasmid and 3.75 µl PLUS reagent (Invitrogen 15338100) following the manufacturer’s instructions. Media was replaced at 24 hours post-transfection and cells were sorted at 36 hours post-transfection using FACS for the top 10% GFP-positive cells. After sorting, cells were seeded to obtain single clones in complete growth media containing 50 µg/ml gentamicin. Knockout clones were validated by Western blotting and genotyping. HT29 / RIPK1-KO clone aA3 was generated using gRNAa, HT29 / RIPK1- KO clone bC5 with gRNAb, HT29 / RIPK1-KO clone cA5 with gRNAc. HCT116/RIPK1-KO clones used in this study were all generated with gRNAc.

### SARS-CoV-2 infection of Calu-3 cells

The Wuhan-like early European SARS-CoV-2 B.1, Freiburg isolate (FR4286, kindly provided by Professor Georg Kochs, University of Freiburg) (Hoffmann *et al*, 2020), was propagated in Vero cells expressing human TMPRSS2 (Olagnier *et al*, 2020) and virus titer determined by TCID_50%_ as previously described (Fougeroux *et al*, 2021). Stocks were validated by sequencing before use in experiments.

Calu-3 epithelial lung cancer cells were cultured in DMEM (Lonza) supplemented with 10% heat inactivated fetal calf serum, 200 IU/mL penicillin, 100 μg/mL streptomycin and 600 μg/mL L-glutamine prior to infection experiments. Calu-3 cells were seeded in flat-bottom 12-well plates (2x10^5^ cells/well in 1 ml media) or flat-bottom 6-well plates (5x10^5^ cells/well in 2 ml media). Upon reaching confluency of 50-70% 24-48 h after seeding, the cells were infected with SARS-CoV2 B.1 at MOIs of 0.5 or 2.0. The plates were incubated at 37°C and tilted every 15 minutes for 1 h to allow for virus adsorption. After 1 h, the media was replaced with fresh media. Supernatant and cell lysates were harvested after 16, 24, 48 and 72h respectively.

To inactivate any live virus prior to biochemical analyses, cell-free supernatant was incubated in 0.5% Triton-X (Sigma Aldrich 11332481001) for 30 minutes at room temperature. Cells for qPCR were lyzed in RNeasy lysis buffer (Qiagen 79254) containing 1% β-Mercaptoethanol and incubated for 10 minutes at room temperature. Cells for Western blotting were lyzed in RIPA buffer (Thermo Fisher 89901), supplemented with with 4x XT sample buffer (BioRad 161-0791) and XT Reducing Agent (BioRad 161-0792), and heated at 95°C for 5 minutes. Virus-inactivated samples were used for subsequent analysis by ELISA, qPCR and Western blotting.

### Transwell migration assays

For transwell migration assays, HT29 / Tet-On-ZBP1 cells were stimulated in FBS-free McCoy’s 5A media supplemented with 0.5% BSA (Sigma A9647, HT29 chemotaxis buffer). Supernatant was collected by centrifuging at 300 x*g* for 5 min or by filtering through 0.2 µm CA syringe filter. THP1 or primary neutrophils were resuspended in FBS-free RPMI media supplemented with 0.5% BSA to an approximate of 1 million/ml, and 0.1 ml were plated into each upper chamber of a 24-well plate (Corning 2421). 0.5 ml conditioned media or HT29 chemotaxis buffer were plated into the lower chamber. The chambers were incubated at 37° C, 5% CO_2_ for 3 hours or as indicated in the figure. Migrated cells were collected from the lower chambers and mixed with Countbright Absolute counting beads (Thermo Fisher C36950) to determine cell density together with the plating population to obtain an accurate plating number. The volume of collected cells was determined by the weight difference of the collection tube before and after collection, assuming density of 1 g/ml. The migration percentage is calculated by dividing migrated cell numbers by plated cell numbers.

### Cell viability assay

The CellTitre-Glo^®^ 2.0 reagent (Promega G9242) was used to determine cell viability according to the manufacturer’s protocol. Cells were seeded in opaque 96-well plates in 100 µl complete growth media at a density of 3.3x10^3^ cells/well for 72-hour time courses or 1.3x10^4^ cells/well for 24-hour treatment. The next day, cells were stimulated with the appropriate chemicals (doxycycline, Sigma, D3072) diluted in 5 µl OptiMEM. Relative viability is calculated by dividing the average of measurement luminescence values of technical replicates for each treatment condition by one reference condition as specified for each figure.

### NF-κB dual-luciferase reporter assay

NF-κB dual luciferase assays were performed with the Dual Luciferase Reporter Assay System kit (Promega E1960). For ZBP1-expression NF-κB reporter assays, cells were plated in 24-well plates at a density of 4x10^5^ cells/well. The next day, they were transfected using per well 125 ng pBIIX-Luc (NF-κB reporter plasmid) (Saksela & Baltimore, 1993), 25 ng SV40- Renilla luciferase plasmid, 0.5 ng ZBP1-expressing pLenti6.3 plasmid (with pBabe-puro or pLenti6.3-GFP as control), and, where indicated, 100 ng pBabe-CYLD plasmids (with pBabe-puro as EV control), 100 ng pcDNA3-OTULIN plasmids (with pcDNA3 as EV control), or 10 ng pcDNA3-GFP-SUB plasmids, in a mixture with 20 μl OptiMEM and 1μl FuGENE HD or FuGENE 6 (Promega E2691) per well. Stimulations were applied immediately after transfection where indicated. Following 24 h incubation, cells were lysed in 75 µl 1x Passive Lysis Buffer (provided in Promega E1960). Luminescence intensity was measured from 10 µl aliquots in technical duplicates using 50 μl of each luciferase assay reagent from the kit.

NF-κB induction levels were calculated by dividing the value of NF-κB luciferase activity by the value of control Renilla luciferase activity. Where indicated, NF-κB induction levels were further normalized to one reference condition.

Expression levels of proteins of interest in the passive lysis buffer lysates were determined by Western blotting as indicated.

### Western blotting

Cells for Western blotting were lysed in 100 µl RIPA buffer (50 mM Tris, pH 7.5, 150 mM NaCl, 1.0% NP-40, 0.5% sodium deoxycholate, 0.1% SDS) supplemented with protease inhibitor (Roche 04693124001) and phosphatase inhibitor cocktails (Roche 4906837001) per well from 6-well plates.

Protein samples were resolved using Bis-Tris gels (10% or 12% acrylamide-bis-acrylamide, 375 mM Tris pH 8.8, 0.1% SDS, 0.1% ammonium persulfate, 0.004% TEMED) in an SDS running buffer (192 mM glycine, 25 mM Tris base, 0.1% SDS) or using NuPAGE^TM^ 4-12% gradient gels (Invitrogen WG1403BX10) in NuPAGE^TM^ MOPS-SDS running buffer (Invitrogen NP0001-02). Separated proteins were transferred to a nitrocellulose membrane (GE 10600002) or PVDF membrane (Millipore IPVH00010). Membranes were blocked for a minimum of 0.5 hour at room temperature in a 5% (w/v) skimmed milk solution in PBST (1.47 mM KH_2_PO_4_, 2.68 mM KCl, 136.9 mM NaCl, 7.97 mM Na_2_HPO_4_, 0.1% Tween-20) with 0.2% sodium azide before incubated in the respective primary antibody overnight at 4°C. Membranes were then washed 3 times for 10 min/time with PBST, incubated with secondary antibody diluted 1:5000-1:10000 in PBST for 1 h, and washed another 3 times for 10 min/time with PBST. Membranes were developed with Amersham^TM^ ECL Western blotting reagent (GE Healthcare RPN2209) with 0-20% Amersham^TM^ ECL Select^TM^ reagent depending on the signal intensity (GE Healthcare RPN2235). Where necessary, membranes were stripped in 1x ReBlot Plus Strong Antibody Stripping Buffer (Millipore 2504) for 15 minutes before blocked again for reblotting.

Primary antibodies used for Western blotting in this study include: anti-ZBP1 (Cell Signaling 60968), anti-Actin, clone C4 (Millipore MAB1501), anti-RIPK3 (Cell Signaling 13526), anti-phospho-p65 (Cell Signaling 3033), anti-p65 (Cell Signaling 8242), anti-phospho-p38 (Cell Signaling 4511), anti-p38 (Cell Signaling 9212), anti-phospho-JNK (BD 612540), anti-JNK (Cell Signaling 9258), anti-phospho-RIPK3 (S227) (Abcam ab209384), anti-RIPK1 (Cell Signaling 3493S), anti-Ubiquitin (rabbit, Cell Signaling, 43124), anti-Ubiquitin (mouse, Cell Signaling 3936), anti-GFP (Cell Signaling 2555), anti-CYLD (Cell Signaling 8462), anti-OTULIN (Cell Signaling 14127), anti-HOIP (R&D AF8039), anti-M1-Ub (kindly provided by David Komander and Rune Busk Damgaard), anti-HOIL-1L (Novus NBP1-88301), anti-SHARPIN (Protein tech 14626-1-AP), anti-TAK1 (Cell Signaling 4505), anti-IKK β (Cell Signaling 8943), anti-K63-Ub (Cell signaling 5621), anti-cIAP1 (Enzo ALX-803-335), anti-FLIP (Cell Signaling 3210), anti-Cleaved caspase-8 (Cell Signaling 9496), anti-Caspase-8 (D35G2) (Cell Signaling 4790) and anti-Caspase-8 (C15) (Adipogen, AG-20B-0057). Secondary antibodies used for Western blotting in this study include: anti-Mouse IgG-HRP (Dako P0447), anti-Rabbit IgG-HRP (Bio-Rad 1706515), anti-Sheep IgG-HRP (R&D HAF016), anti-human IgG-HRP (Bio-Rad 172-1033) and anti-Rat IgG-HRP (Thermo Fisher 31470).

### Anti-FLAG immunoprecipitation

To perform anti-FLAG immunoprecipitation for ZBP1-associated proteins, cells were plated in 10 cm dishes at a density of 3-4x10^6^ cells/dish, 3-6 dishes/condition, and stimulated with 500 ng/ml Dox the next day for 16 h. After stimulation, cells were washed twice with PBS and lysed in 500 µl/dish TBSN buffer (50 mM Tris-HCl, pH 7.5, 150 mM NaCl, 0.5% NP-40) supplemented with protease inhibitor cocktail, phosphatase inhibitor cocktail and 50 mM N-Ethylmaleimide (NEM; Sigma Aldrich E1271) by ice incubation for a minimum of 15min. Lysates were centrifuged at maximum speed for 10 minutes at 4°C to remove debris pellets. After taking input samples, the lysate supernatant was precleared with 10 µl/dish mouse IgG-Agarose beads (Sigma A0919) for a minimum of 30min at 4°C with end-over-end rotation, before incubated with 10 µl/dish anti-FLAG M2 agarose beads (Sigma A2220) for a minimum of two hours with end-over-end rotation at 4°C. After enrichment, beads were washed five times with 1 ml TBSN buffer and eluted in 13.3 µl/dish 2xLSB by shaking at 95°C, 750rpm, for 10min.

Western blotting was used to probe for co-immunoprecipitated proteins. Equivalent of eluate materials from one 10 cm dish was loaded on each pull-down lane, and 1% input was loaded for reference except for ZBP1, for which 5% input was loaded.

### Enrichment of Ub-conjugates

To enrich for ubiquitinated proteins, cells were seeded in 10 cm dishes and stimulated as described for FLAG immunoprecipitation. For GST-1xUBA and SUB pulldown, beads were prepared by incubation with recombinant GST-1xUBA or SUB for at least 1h at 4°C with rotation and washed three times with TUBE lysis buffer (20 mM sodium phosphate buffer, pH 7.4, 1% NP-40, 2 mM EDTA). For GST-1xUBA, 10µl/dish Glutathione Magnetic Agarose beads (Thermo Fisher 78602) were incubated with 30 µg/dish GST-1xUBA. K63-SUB-bound beads were prepared by incubating 20 µl/dish Streptavidin Magnetic Agarose beads (Thermo Fisher 88817) with 5 µg/dish recombinant biotinylated K63-SUB. For M1-SUB pulldown, 10 µl/dish Glutathione Magnetic Agarose beads were incubated with 50 µg M1-SUB (Fiil *et al*, 2013).

Cells were lysed using 500 µl TUBE lysis buffer supplemented with protease inhibitor cocktail, phosphatase inhibitor cocktail, 50 mM NEM and 1 mM DTT following the same protocol as for FLAG immunoprecipitation. After taking input samples, centrifuged lysate supernatant was incubated with 10 µl/dish GST-1xUBA- or SUB-bound beads overnight at 4°C with end-over-end rotation. After enrichment, beads were washed three times with TUBE lysis buffer and heated in 2xLSB for 10 minutes at 95°C for elution.

Equivalent of pull-down samples from one 10 cm dish and 1% input (except for ubiquitin blot, for which 5% input was loaded) were resolved on 4-12% NuPAGE gradient gels in NuPAGE MOPS-SDS running buffer and transferred onto PVDF membrane for Western blotting.

### On-bead deubiquitinase treatment

On-bead deubiquitinase digestion was performed to probe for the existence and composition of ubiquitin chains in FLAG-immunoprecipitated or TUBE-enriched sections. After enrichment, beads were washed three times with lysis buffer and incubated shaking with or without 1 µM USP21 (R&D E-622-050) in 30 µl DUB buffer (20 mM HEPES, pH 7.5, 100 mM NaCl, 1 mM MnCl_2_, 0.01% w/v Brij-35, 5 mM DTT) for 1h at 30°C before LSB buffer was added to end the reaction.

### Enzyme-linked immunosorbent assay (ELISA)

For ELISAs, cells were plated at a density of 1x10^5^ per well in 24-well plates or 2x10^5^ per well in 12-well plates, and stimulated the next day for 24h with the appropriate chemicals (doxycycline, Sigma D3072; dimerizer, Clontech 635059; (5Z)-7-Oxozeaenol / TAK1i, Tocris 3604; IKK inhibitor VII, Merck Millipore 401486; IKK inhibitor XII, Merck Millipore 401491; zVAD, Santa Cruz sc-311560; Nec1s, Enzo BV-2535-1; GSK’872, Merck Millipore 5.30389.001; NSA, R&D 5025/10). Cell culture supernatants were centrifuged at 300 x*g* for 5 minutes to remove debris, diluted as appropriate, and loaded in technical duplicates or triplicates. Measurement of cytokine concentrations was carried out with R&D DuoSet ELISA kits (CXCL8, DY208-05, CXCL1, DY275-05, CXCL10, DY266-05, IL-6, DY206-05) according to manufacturer’s instructions. For data analysis, the absorbance at 540nm of each well was subtracted from the 450nm value. Generation of standard curves and interpolation of data were performed in GraphPad Prism. Where indicated, cells were collected after stimulation for Western blotting to assess the protein expression levels.

### siRNA-mediated gene knockdown

Knockdown of RIPK3 was achieved using Accell siRNA SMARTpool (Horizon Discovery E- 003534-00-0005, Gene ID 11035), with Accell siRNA non-targeting pool (Horizon Discovery D-001910-10-05) as control targeting mismatch sequence (siMM). Cells were plated at 40% confluency. The next day, siRNA was transfected using Dharmafect transfection reagent (Horizon Discovery T-2001-03) in a mixture of 200 µl OptiMEM (Gibco 31985), 3 µl Dharmafect and 0.03 nmol siRNA per 200,000 cells. 24h after transfection, cells were split into desired number and format of wells in complete growth medium for treatment.

Knockdown of HOIP was achieved using custom-synthesized siRNA from Sigma (sense strand: GGCGUGGUGUCAAGUUUAA[dT][dT]; antisense strand: UUAAACUUGACACCACGCC[dT][dT]) (Haas *et al*, 2009)). siRNA against mismatch sequence (Sigma SIC001) was used as non-targeting control. Cells were plated in 12-well plates at 0.8-1.6x10^5^ per well, and transfected with 0.04 nmol siRNA in a mixture with 100 µl OptiMEM and 1 µl Lipofectamine RNAiMAX reagent (Invitrogen 100014472). 24h after siRNA transfection, cells were trypsinized and reseeded for indicated applications.

### RNA isolation, cDNA synthesis and qPcR

RNeasy Mini kit (Qiagen 79254) was used for RNA isolation according to the manufacturer’s instructions. and on-column DNA digestion was performed with RNase-free DNase Set (Qiagen 74106). For cDNA synthesis, around 1 µg RNA (10 µl) was incubated with 1 µl 10 µM random pentadecamers (IDT 169190224) and 0.5 µl 100µM anchored oligo(dT)_20_ primer (Sigma, custom synthesized) at 65°C for 5 minutes, before supplemented with 0.5 µl RevertAid reverse transcriptase (Thermo Scientific EP0441) and 0.5 µl RiboLock RNase inhibitor (Thermo Scientific EO0381) in a total volume of 20 µl RevertAid Reverse Transcriptase buffer (Thermo Scientific LT-02241). Samples were then subject to reverse transcription program of 10 minutes at 25°C, followed by 60 minutes at 42°C and 10 minutes at 70°C. The resulting cDNA was diluted as appropriate before quantitative PCR.

Quantitative PCR from the cDNA was performed using the SYBR Select qPCR mastermix (Applied Biosystems 4472908) using 2 μl cDNA, 1 µM forward primer and 1 µM reverse primer per reaction in a 10 µl reaction volume in technical duplicates. Primer pairs used were as following:

Hypoxanthine phosphoribosyltransferase (*HPRT*; used as reference for normalization), 5’- AGCCAGACTTTGTTGGATTTG-3’ and 5’-TTTACTGGCGATGTCAATAGG-3’; *CXCL8*, 5’- TCTGGCAACCCTAGTCTGCT-3’ and 5’-AAACCAAGGCACAGTGGAAC-3; *CXCL1*, 5’-TCCTGCATCCCCCATAGTTA-3’ and 5’-CTTCAGGAACAGCCACCAGT-3’; *CXCL2*, 5’-GTGACGGCAGGGAAATGTAT-3’ and 5’- CACAGAGGGAAACACTGCAT-3’; *CXCL3*, 5’- AGGGACAGCTGGAAAGGACT-3’ and 5’- AAGGTGGCTGACACATTATGG-3’; *ZBP1,* 5’-GAAGCAAGAATTCCCAGTCCAG-3’ and 5’-TCGAGAAAGCACGATTTCATGT-3’; *TNF*, 5’- TGCTGCAGGACTTGAGAAGA-3’ and 5’-GAGGAAGGCCTAAGGTCCAC-3’; *CXCL10*, 5’-GTGGATGTTCTGACCCTGCT-3’ and 5’- GAGGATGGCAGTGGAAGTCC-3’;

Interpolation of CT values was performed with the built-in software of the qPCR machine (Roche LightCycler 480 or Applied Biosystems 7500). Relative mRNA expression levels were calculated with the comparative CT method (Schmittgen & Livak, 2008).

## QUANTIFICATION AND STATISTICAL ANALYSIS

### SARS-CoV-2 single cell RNA-seq expression analysis

Processed single cell RNA TPM expression of (Ren *et al*, 2021) associated with GSE158055 were downloaded and analysed from http://covid19.cancer-pku.cn/#/summary. The patient TPM expression of a gene of interest was computed as the mean TPM expression of the patients’ individual constituent single cells.

### Statistical analysis for quantitative experiments

Statistical analyses were carried out in GraphPad Prism. The statistical details are described in figure legends, with n representing the number of biological replicates. Outliers were identified by ROUT method (Q=1%) and excluded from statistical analyses.

For the comparison between two conditions, an F-test was used to test for the statistical differences between the standard deviations (SDs) of the two conditions. A two-tailed unpaired t-test was used when SDs from the two conditions were not significantly different. A Welch’s t-test was used if SDs from the two samples are significantly different.

For comparison of more than two conditions, a Brown-Forsythe test was used to test for statistical differences between the SDs of all conditions. When the Brown-Forsythe test found no significant differences between group variances, one-way ANOVA and Sidak’s multiple comparisons test were used to test for statistical differences between indicated conditions.

When the Brown-Forsythe test found a significant difference between group variances, Brown-Forsythe and Welch ANOVA tests and Dunnet’s T3 multiple comparisons test were used. Multiple t-tests were used when SDs from conditions were found to be significantly different and Brown-Forsythe and Welch ANOVA tests were not possible.

For grouped data or data from multi-factor experiment designs, two-way ANOVA and the following multiple comparison tests were used following Prism’s recommendation on the choice of multiple comparison tests.

The distribution of the single cell RNA sequencing datasets were found significantly different from Gaussian distribution, and therefore they were analyzed using Kruskai-Wallis test and Dunn’s multiple comparisons test.

## Acknowledgements

This work was supported by the Ludwig Institute for Cancer Research Ltd. Work in the M.G-H. llab was supported by a Wellcome Trust Fellowship (102894/Z/13/Z and 215612/Z/19/Z) and the LEO foundation. R.P. was supported by a China Scholarship Council-University of Oxford DPhil Scholarship (2015-2019; GAF1516_CSCUO_841725). S.R.P. was funded by The Independent Research Fund Denmark (0214-00001B) the European Research Council (ERC-AdG ENVISION; 786602), and the Novo Nordisk Foundation (NNF18OC0030274). We acknowledge the contribution to this study made by the Oxford Centre for Histopathology Research and the Oxford Radcliffe Biobank, which are supported by the University of Oxford, the Oxford CRUK Cancer Centre and the NIHR Oxford Biomedical Research Centre (Molecular Diagnostics Theme/Multimodal Pathology Subtheme), and the NIHR CRN Thames Valley network. We thank Prof. David Komander and Prof. Rune Busk Damgaard for providing Met1-Ubiquitin antibody, Dr. Norbert Volkmar and Prof. John Christianson for providing shRNA plasmids targeting mismatch sequences, Tetralogic Pharmaceuticals for Compound A, Professor Georg Kochs, for providing SARS-CoV-2 virus stock, Dr. Amit Shrestha and Prof. Nick La Thangue for providing HL60 cells, and members of the M.G-H. group for helpful suggestions and reading the manuscript.

## Author Contributions

M.G-H. and R.P. conceptualized the project. R.P. designed and performed most of the experiments. X.W-K. performed experiments in Figures 1B, 1D, 4H, 5A, 5C, 5D, S1A, S1D-F and technically assisted other experiments. M.I. performed virus infection assays for Figures 7C, 7D, 7F, 7G and S7B-D. F.Y.Z. helped with data analysis in Figure 7B, 7E, S3B and S7A. J.McCarthy helped with the access to and preparation of primary donor samples under the supervision of C.S.L.. S.L.O., A.O., J.R. and J.Maelfait provided unpublished reagents and advised on experimental design. M.G-H., S.R.P. and X.L. acquired funds. M.G-H. supervised the work. K.B. co-supervised the work at early stages. R.P. and M.G-H. wrote the manuscript. X.W-K., K.B., S.R.P., J.Maelfait, F.Y.Z., M.I. and J.McCarthy reviewed the manuscript.

