## Supplemental Information for "ZBP1 induces inflammatory signaling via RIPK3 and promotes SARS-CoV-2-induced cytokine expression"

**Figure S1**

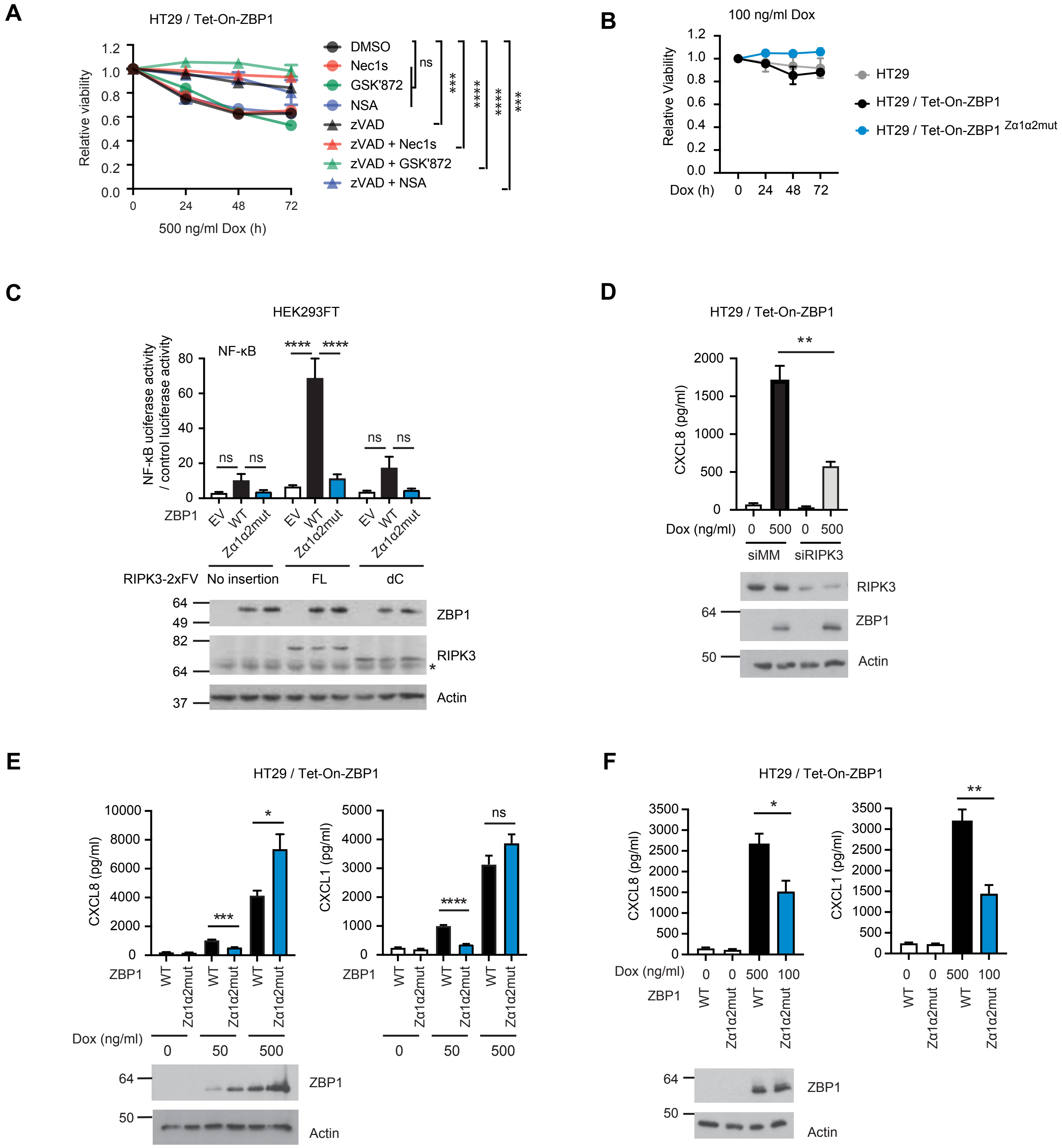
A
HT29 / Tet-On-ZBP1
Relative viability
500 ng/ml Dox (h)
DMSO
Nec1s
GSK'872
NSA
zVAD
zVAD + Nec1s
zVAD + GSK'872
zVAD + NSA
ns
***
****
****
***
B
100 ng/ml Dox
Relative Viability
Dox (h)
HT29
HT29 / Tet-On-ZBP1
HT29 / Tet-On-ZBP1 Zα1α2mut
C
HEK293FT
NF-κB
NF-κB uciferase activity / control luciferase activity
ZBP1
EV
WT
Zα1α2mut
RIPK3-2xFV
No insertion
FL
dC
ZBP1
RIPK3
Actin
D
HT29 / Tet-On-ZBP1
CXCL8 (pg/ml)
Dox (ng/ml)
siMM
siRIPK3
RIPK3
ZBP1
Actin
E
HT29 / Tet-On-ZBP1
CXCL8 (pg/ml)
CXCL1 (pg/ml)
ZBP1
Dox (ng/ml)
ZBP1
Actin
F
HT29 / Tet-On-ZBP1
CXCL8 (pg/ml)
CXCL1 (pg/ml)
Dox (ng/ml)
ZBP1
ZBP1
Actin

**Figure S1. Related to Figure 1. ZBP1 expression induces RIPK3-mediated NF-κB activation independently of cell death.**

(A) Relative viability levels of HT29 / Tet-On-ZBP1 cells treated with 500 ng/ml Dox for up to 3 days in combination with 20 μM zVAD, 10 μM Nec1s, 10 μM GSK'872 and/or 1 μM NSA as indicated. Values are normalized to untreated wells. Data is plotted as mean with S.E.M. (n = 4). Two-way ANOVA and Dunnet's multiple comparisons tests were used to test for statistical differences between indicated conditions. ns, $p > 0.99$, ***, $p = 0.0006$ for zVAD, $p = 0.0007$ for zVAD+NSA, ****, $p<0.0001$.

(B) Relative viability of cells treated with 100 ng/ml Dox for up to 3 days as indicated. Cell viability at the end of treatment was determined by measuring ATP levels. Measurement values were normalized to that of 0 h within each cell line. Data is presented as mean with S.E.M (n = 3).

(C) Folds of NF-κB activity over control luciferase in HEK293FT cells stably expressing variants of RIPK3 and transiently transfected with variants of ZBP1, co-expressing dual luciferase reporters. Reporter activities were measured 24 h after transfection. Data is presented as mean with S.E.M (n = 3). One-way ANOVA and Sidak's multiple comparisons test were used to test for statistical differences between indicated conditions. ns, $p > 0.2$, ****, $p<0.0001$. Cell lysates were loaded for western blot analysis. Asterisk indicates background signals of the antibody.

(D) CXCL8 concentrations in cell culture supernatants of HT29 / Tet-On-ZBP1 cells transfected with siRNAs targeting mismatch sequence (MM) or RIPK3 and treated with 500 ng/ml Dox for 24 h. Data is presented as mean with S.E.M. (n = 3). Unpaired t-tests were used to test for statistical differences between indicated conditions. **, $p = 0.0052$. Cells were lysed for western blot.

(E. F) Chemokine concentrations in cell culture supernatants of HT29 cells induced to express WT or Zα1α2mut ZBP1 for 24 h with indicated Dox concentrations. Cells from the same wells were lysed for western blot to determine ZBP1 expression levels. (E) Data is presented as mean with S.E.M (n = 6). Brown-Forsythe and Welch ANOVA tests and Dunnet's T3 multiple comparisons test were used to test for statistical significances between indicated conditions. ***, p = 0.0002, *, p = 0.0447, ****, p<0.0001, ns, p = 0.2153. (F) Data is presented as mean with S.E.M (n = 4). Unpaired t-tests were used to test for the statistical differences between the indicated conditions. *, p = 0.0257, **, p = 0.0034.

### Figure S2

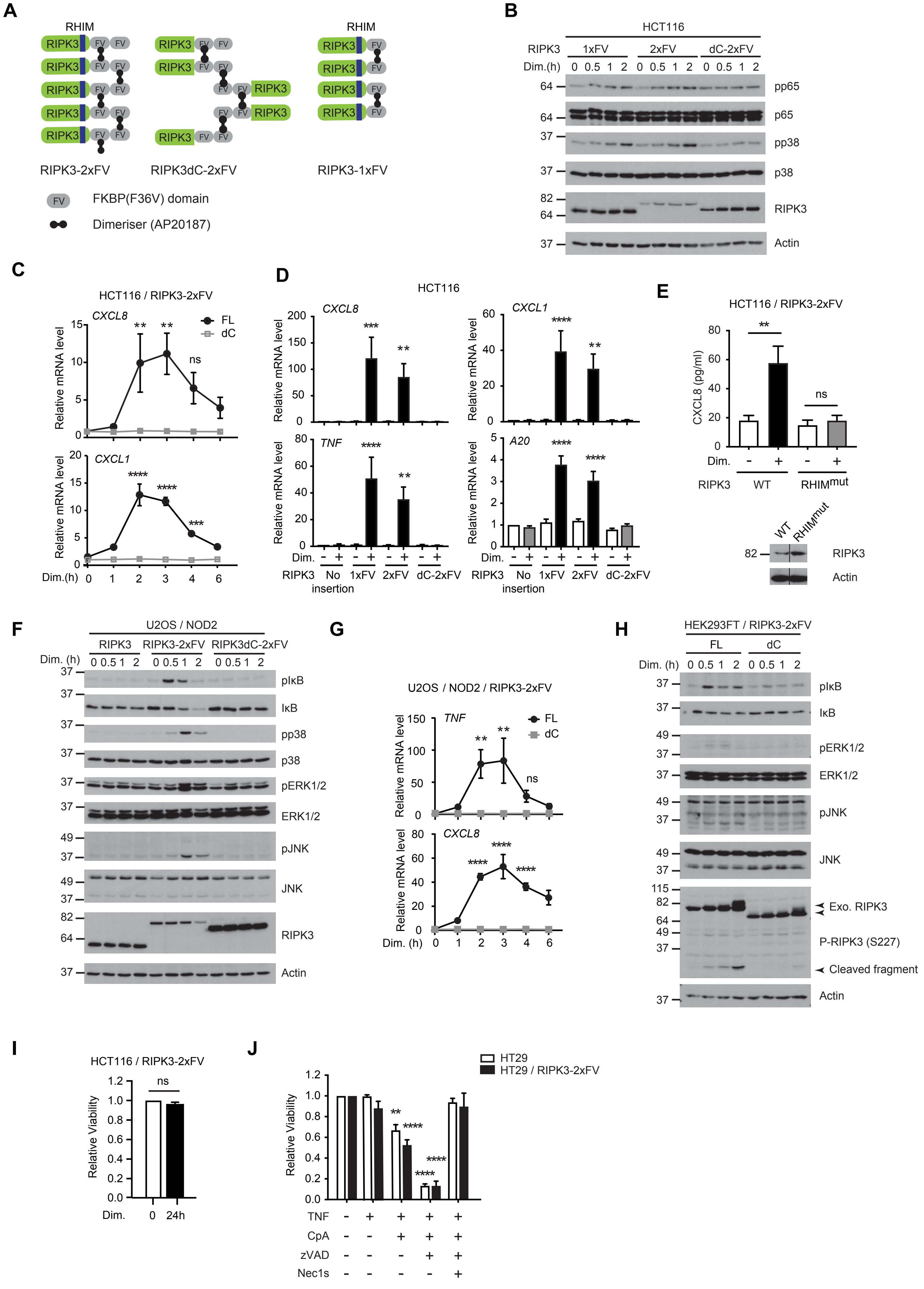
A
RHIM
RIPK3
FV
RIPK3-2xFV
RIPK3dC-2xFV
RIPK3-1xFV
FKBP(F36V) domain
Dimeriser (AP20187)
B
HCT116
RIPK3 1xFV 2xFV dC-2xFV
Dim.(h) 0 0.5 1 2 0 0.5 1 2 0 0.5 1 2
pp65
p65
pp38
p38
RIPK3
Actin
C
HCT116 / RIPK3-2xFV
CXCL8
CXCL1
FL
dC
Relative mRNA level
Dim.(h) 0 1 2 3 4 6
D
HCT116
CXCL8
CXCL1
TNF
A20
Relative mRNA level
Dim.
RIPK3 No insertion 1xFV 2xFV dC-2xFV
E
HCT116 / RIPK3-2xFV
CXCL8 (pg/ml)
Dim.
RIPK3 WT RHIMmut
WT
RHIMmut
RIPK3
Actin
F
U2OS / NOD2
RIPK3 RIPK3-2xFV RIPK3dC-2xFV
Dim. (h) 0 0.5 1 2 0 0.5 1 2 0 0.5 1 2
pIκB
IκB
pp38
p38
pERK1/2
ERK1/2
pJNK
JNK
RIPK3
Actin
G
U2OS / NOD2 / RIPK3-2xFV
TNF
CXCL8
FL
dC
Relative mRNA level
Dim. (h) 0 1 2 3 4 6
H
HEK293FT / RIPK3-2xFV
FL dC
Dim. (h) 0 0.5 1 2 0 0.5 1 2
pIκB
IκB
pERK1/2
ERK1/2
pJNK
JNK
Exo. RIPK3
P-RIPK3 (S227)
Cleaved fragment
Actin
I
HCT116 / RIPK3-2xFV
Relative Viability
Dim. 0 24h
J
HT29
HT29 / RIPK3-2xFV
Relative Viability
TNF
CpA
zVAD
Nec1s

**Figure S2. Related to Figure 1. Chemical-induced RIPK3 oligomerization drives inflammatory response.**

(A) Schematic illustration of chemical-inducible RIPK3 oligomerization system. The RIPK3dC-2xFV molecules may not be able to align in stacks due to lack of the RHIM domain.

(B) Western blot analysis of HCT116 cells stably expressing FV-fused RIPK3 variants treated with 100 nM dimerizer.

(C) Time courses of mRNA levels of *CXCL8* and *CXCL1* in HCT116 cells stably expressing RIPK3-2xFV (FL) or RIPK3dC-2xFV (dC) in response to treatment with 100 nM dimerizer. Data is plotted as mean with S.E.M. (n = 3). Two-way ANOVA and Tukey's multiple comparisons test were used to test for statistical differences between 0 h and indicated time points in FL cells. ns, p = 0.1360, **, p = 0.0046 for FL 2 h, p = 0.0012 for FL 3 h, ****, p < 0.0001, ***, p = 0.0008.

(D) Relative mRNA levels of representative NF-κB target genes in HCT116 cells stably expressing FV-fused RIPK3 variants treated with 0 or 100 nM dimerizer for 3 h. Data is presented as mean with S.E.M. (n = 4). Two-way ANOVA and Sidak's multiple comparison tests were used to test for statistical differences between indicated condition and solvent-treated condition within each cell line. ***, p = 0.0001, **, p = 0.0060 for *CXCL8*, p = 0.0018 for *CXCL1*, p = 0.0038 for *TNF*, ****, p < 0.0001.

(E) CXCL8 concentrations in cell culture supernatants of HCT116 cells stably expressing WT or RHIM-mutant ($RHIM^{mut}$) RIPK3-2xFV treated with 100 nM dimerizer for 24 h. $RHIM^{mut}$ RIPK3 carries mutation in the three key residues, VQV, of the RHIM region, to tetra-alanine (AAAA). Data is plotted as mean with S.E.M. (n = 5). One-way ANOVA and Sidak's multiple comparisons tests were used to test for statistical differences between indicated time points. **, p = 0.0011, ns, p = 0.9301. Cells were lysed for western blot to determine expression levels.

Line indicates that image was cut and spliced to remove non-relevant lanes from the scanned blot.

(F) Western blot analysis of U2OS / NOD2 cells stably expressing RIPK3 variants treated with 100 nM dimerizer.

(G) Time course of relative mRNA levels of *CXCL8* and *TNF* in U2OS / NOD2 cells stably expressing RIPK3-2xFV (FL) or RIPK3dC-2xFV (dC) treated with 100 nM dimerizer. Data is plotted as mean with S.E.M. (n = 3). Two-way ANOVA and Tukey's multiple comparisons test were used to test for statistical differences between 0h and indicated time points in FL cells. **, p = 0.0021 for 2 h, p = 0.0010 for 3 h, ns, p = 0.6323, ****, p < 0.0001.

(H) Western blot analysis of HEK293FT cells stably expressing RIPK3-2xFV or RIPK3dC-2xFV treated with 100 nM dimerizer.

(I) Relative viability of HCT116 / RIPK3-2xFV cells treated with 0 or 100 nM dimerizer for 24 h. Data is presented as mean with S.E.M. (n = 3). A Welch's t-test was used to test for statistical differences between indicated conditions. ns, p = 0.2294.

(J) Relative viability levels of HT29 or HT29 / RIPK3-2xFV cells pretreated with combinations of 100 nM CpA, 20 μM zVAD and 10 μM Nec1s as indicated for 1 h, and then stimulated with 0 or 2 ng/ml TNF for 24h. Measurement values were normalized to DMSO-treated condition within each cell line. Data is presented as mean with S.E.M. (n = 3). Two-way ANOVA and Sidak's multiple comparison test were used to test for statistical differences between indicated condtions and DMSO-treated condition of each cell line. **, p = 0.0044, ****, p < 0.0001.

### Figure S3

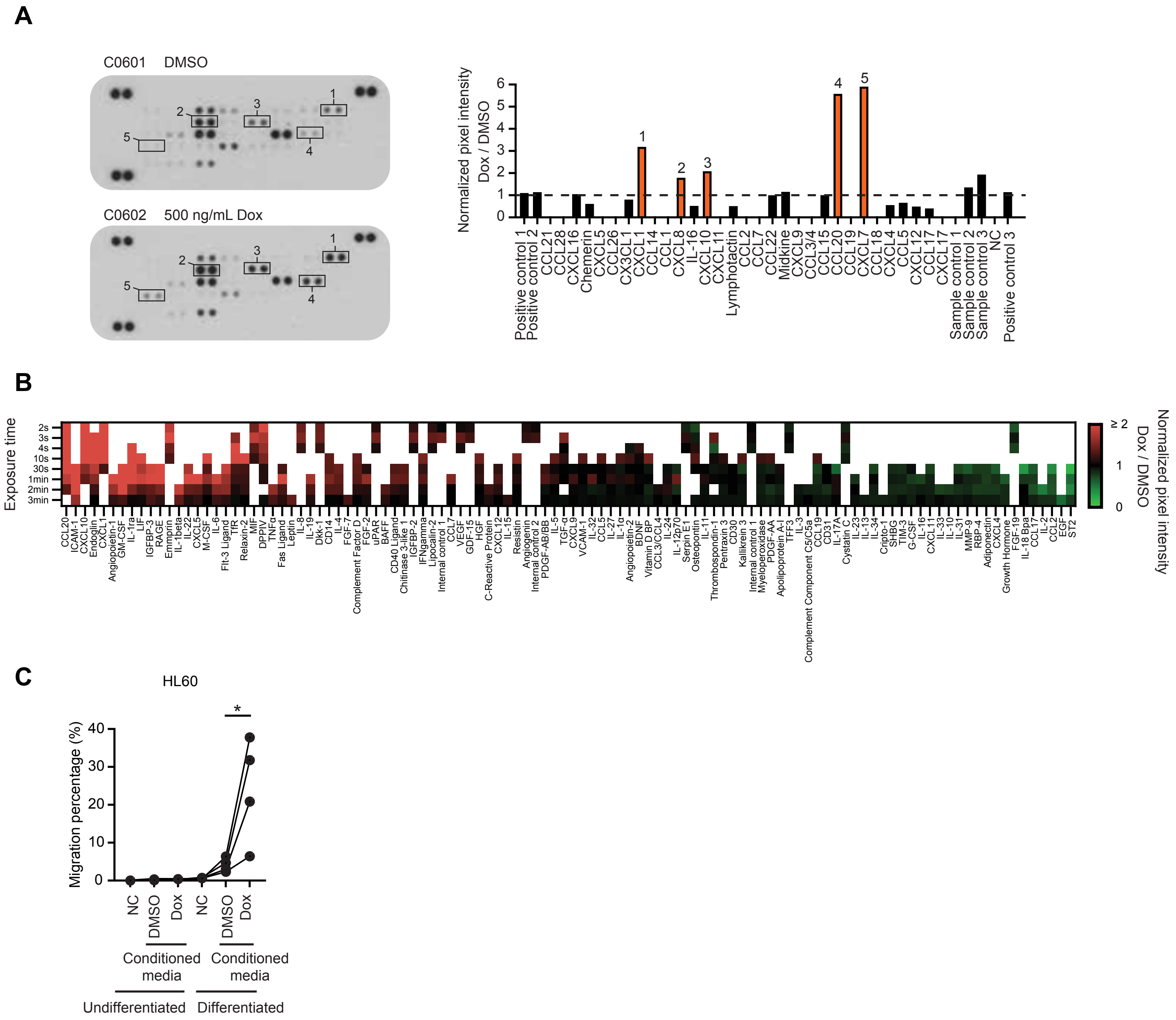
A
C0601 DMSO
C0602 500 ng/mL Dox
1
2
3
4
5
Normalized pixel intensity
Dox / DMSO
0
1
2
3
4
5
6
Positive control 1
Positive control 2
CCL21
CCL28
CXCL16
Chemerin
CXCL5
CCL26
CX3CL1
CXCL1
CCL14
CCL1
CXCL8
IL-16
CXCL10
CXCL11
Lymphotactin
CCL2
CCL7
CCL22
Midkine
CXCL9
CCL3/4
CCL15
CCL20
CCL19
CXCL7
CCL18
CXCL4
CCL5
CXCL12
CCL17
CXCL17
Sample control 1
Sample control 2
Sample control 3
NC
Positive control 3
B
Exposure time
2s
3s
4s
10s
30s
1min
2min
3min
≥ 2
1
0
Normalized pixel intensity
Dox / DMSO
C
HL60
Migration percentage (%)
0
10
20
30
40
*
NC
DMSO
Dox
NC
DMSO
Dox
Conditioned media
Conditioned media
Undifferentiated
Differentiated

**Figure S3. Related to Figure 1. ZBP1 expression induces a proinflammatory secretome.** (A-B) Chemokine (A) and cytokine (B) arrays of the conditioned media from HT29 / Tet-On-ZBP1 cells treated with DMSO or 500 ng/ml Dox for 24 h. Normalized pixel intensity folds of Dox over DMSO from the blots was plotted (n = 1).

(C) Transwell migration percentages of differentiated or undifferentiated HL60 cells towards chemotaxis buffer (NC) or conditioned media from HT29 / Tet-On-ZBP1 cells treated with DMSO or 500 ng/ml Dox for 24 h. Data from each biological replicate of independently differentiated HL60 cells and conditioned media are connected with solid lines (n = 4). A paired t-test were used to determine the statistical difference between indicated conditions. *, p = 0.0447.

### Figure S4

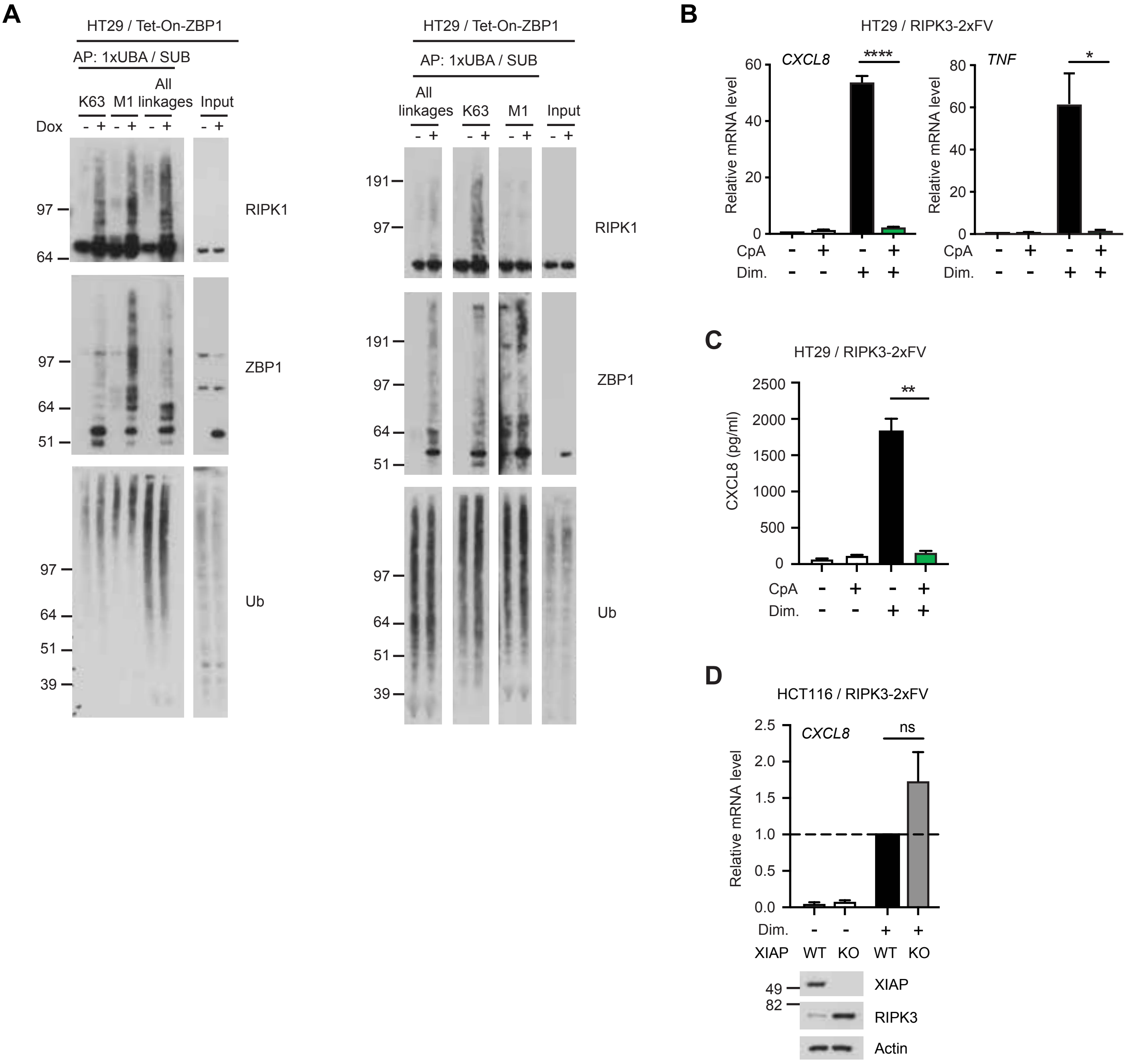

A
HT29 / Tet-On-ZBP1
AP: 1xUBA / SUB
K63
M1
All linkages
Input
Dox
RIPK1
ZBP1
Ub
All linkages
K63
M1
Input
B
HT29 / RIPK3-2xFV
CXCL8
TNF
Relative mRNA level
****
*
CpA
Dim.
C
HT29 / RIPK3-2xFV
CXCL8 (pg/ml)
**
CpA
Dim.
D
HCT116 / RIPK3-2xFV
CXCL8
ns
Relative mRNA level
Dim.
XIAP
WT
KO
WT
KO
XIAP
RIPK3
Actin

**Figure S4. Related to Figure 3 and Figure 4. K63-Ub and M1-Ub facilitate ZBP1 inflammatory signaling**

(A) Two independent biological replicates of Ub-conjugate pull down analyses as in Figure 3C.

(B) Relative mRNA levels of *CXCL8* and *TNF* in HT29 / RIPK3-2xFV cells pretreated with 0 or 2 μM CpA for 1 h before treated with 0 or 100 nM dimerizer for 3 h. Data is plotted as mean with S.E.M. ($n = 3$). Unpaired t-tests were used to test for statistical differences between indicated conditions. ****, $p < 0.0001$, *, $p = 0.0145$.

(C) CXCL8 concentrations in the cell culture supernatant of HT29 / RIPK3-2xFV cells treated with 0 or 100 nM CpA for 1h before treatment with 0 or 100 nM dimerizer for 24 h. Data is plotted as mean with S.E.M. ($n = 3$). A Welch's t-test was used to test for statistical differences between indicated conditions. **, $p = 0.0097$.

(D) Relative *CXCL8* mRNA levels in WT or XIAP-knockout HCT116 cells stably expressing RIPK3-2xFV, treated with 0 or 100 nM dimerizer for 3 h. Data is plotted as mean with S.E.M. ($n = 3$). A Welch's t-test were used to test for statistical differences between indicated conditions. ns, $p = 0.2057$. Cells were lyzed for Western blot analysis of expression levels.

#### Figure S5

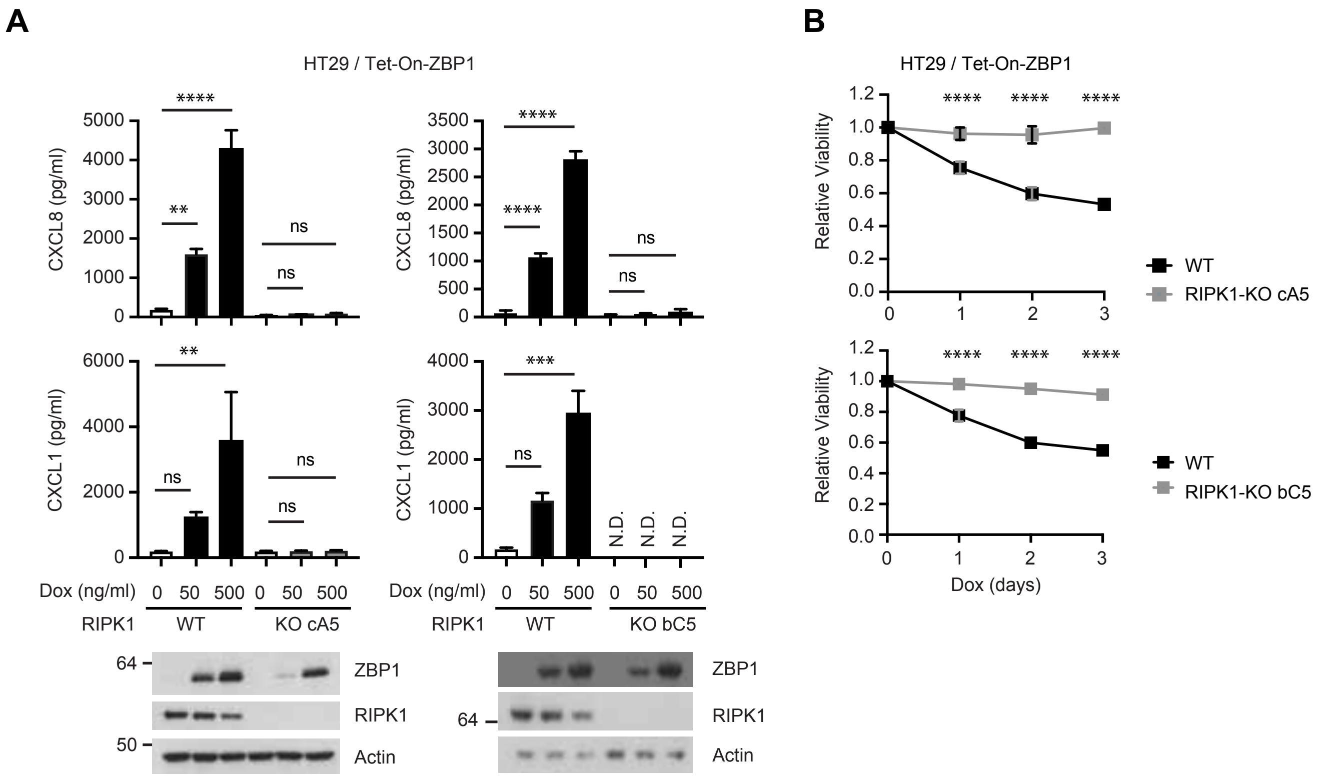

A
B
HT29 / Tet-On-ZBP1
CXCL8 (pg/ml)
CXCL1 (pg/ml)
Dox (ng/ml)
0
50
500
RIPK1
WT
KO cA5
KO bC5
N.D.
ns
64
50
ZBP1
RIPK1
Actin
Relative Viability
Dox (days)
WT
RIPK1-KO cA5
RIPK1-KO bC5

**Figure S5. Related to Figure 5. RIPK1 is required for ZBP1 signaling**

(A) Chemokine concentrations in the supernatants of HT29 / Tet-On-ZBP1 or two independent clones of HT29 / RIPK1-KO / Tet-On-ZBP1 cells treated with 0, 50 or 500 ng/ml Dox. N.D., not detected. data is plotted as mean with S.E.M. ($n = 3$). One-way ANOVA and Sidak's multiple comparisons test were used to test for statistical differences between indicated conditions. ns, $p > 0.05$, **, $p = 0.0079$ for CXCL1, $p = 0.0012$ for CXCL8, ***, $p = 0.0008$, ****, $p<0.0001$. Cell lysates were analyzed by western blot for ZBP1 expression levels.

(B) Relative viability levels of HT29 / Tet-On-ZBP1 cells or two independent clones of HT29 / RIPK1-KO / Tet-On-ZBP1 cells after treated with 500 ng/ml Dox for up to 3 days as indicated. Measurement values are normalized to no treatment within the same cell line. Data is plotted as mean with S.E.M. ($n = 3$). Two-way ANOVA and Sidak's multiple comparison tests were used to test for statistical differences between WT and RIPK1-KO cells one to three days after Dox treatment. ****, $p<0.0001$.

### Figure S6

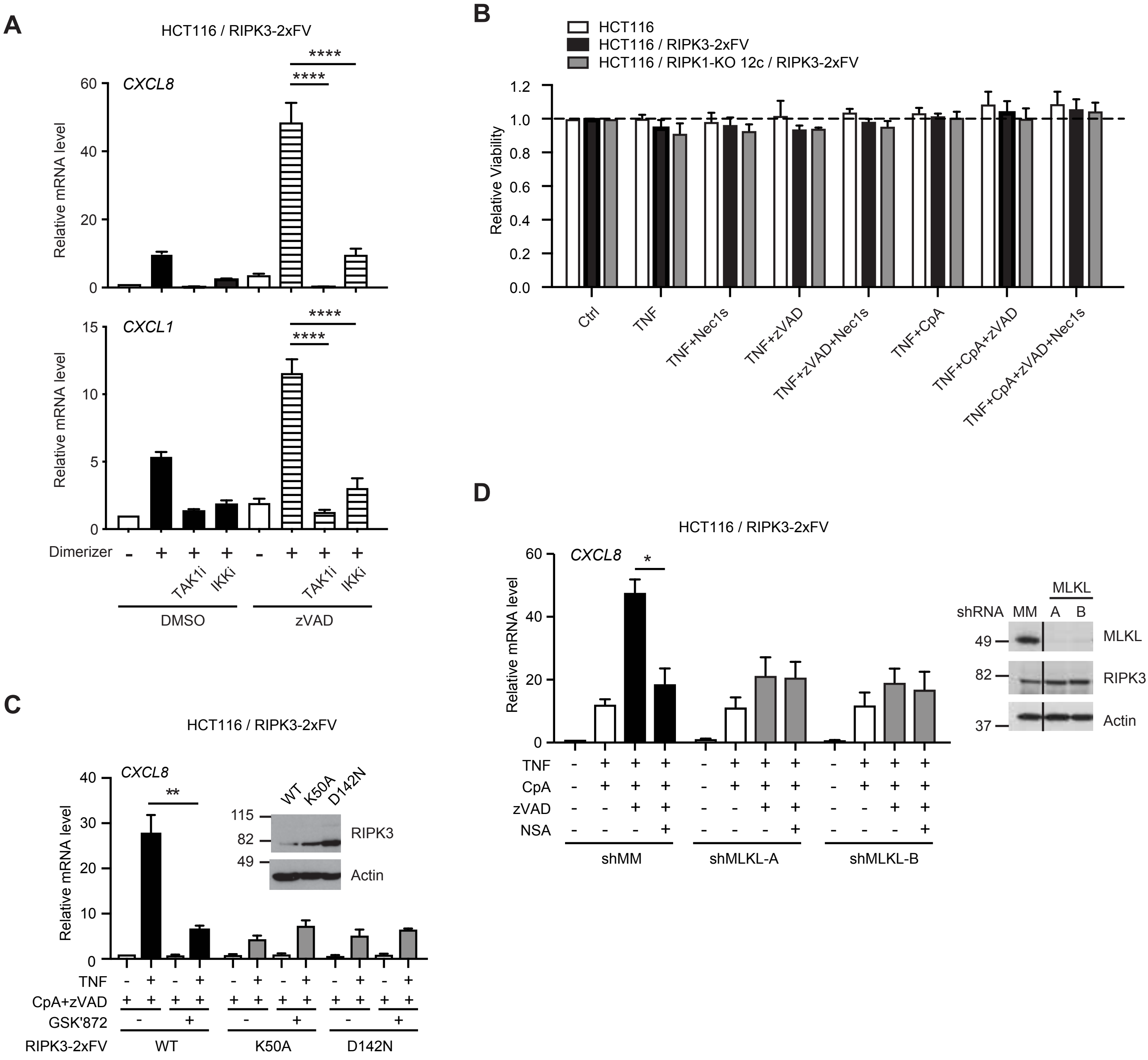
A
HCT116 / RIPK3-2xFV
CXCL8
CXCL1
Relative mRNA level
****
Dimerizer
TAK1i
IKKi
DMSO
zVAD
B
HCT116
HCT116 / RIPK3-2xFV
HCT116 / RIPK1-KO 12c / RIPK3-2xFV
Relative Viability
Ctrl
TNF
TNF+Nec1s
TNF+zVAD
TNF+zVAD+Nec1s
TNF+CpA
TNF+CpA+zVAD
TNF+CpA+zVAD+Nec1s
C
HCT116 / RIPK3-2xFV
CXCL8
Relative mRNA level
**
WT
K50A
D142N
RIPK3
Actin
TNF
CpA+zVAD
GSK'872
RIPK3-2xFV
D
HCT116 / RIPK3-2xFV
CXCL8
Relative mRNA level
*
TNF
CpA
zVAD
NSA
shMM
shMLKL-A
shMLKL-B
MLKL
shRNA
MM
A
B
MLKL
RIPK3
Actin

**Figure S6. Related to Figure 6. Caspase inhibition leads to the activation of RIPK3 kinase activity-dependent inflammatory signaling**

(A) Relative *CXCL8* mRNA levels in HCT116 / RIPK3-2xFV cells pretreated with 0 or 20 μM zVAD in combination with TAK1 or IKK inhibitors for 1 h, before treated with 0 or 100 nM dimerizer for 3 h. Data is presented as mean with S.E.M. (n = 3). One-way ANOVA and Sidak's multiple comparisons test were used to test for the statistical difference between indicated conditions. ****, p < 0.0001.

(B) Relative viability of indicated cell lines treated in the same way as described in Figure 6D. Values were normalized to the DMSO-treated condition of each cell line. Data is presented as mean with S.E.M. (n = 3).

(C) Relative *CXCL8* mRNA levels in HCT116 cells stably expressing WT or catalytic-inactive mutants of RIPK3-2xFV, pretreated with 2 μM CpA, 20 μM zVAD and/or 10 μM GSK'872 as indicated for 1 h, followed by stimulation with 0 or 2 ng/ml TNF for 3 h. Data is presented as mean with S.E.M. (n = 3). An unpaired t-test was used to test for the statistical difference between indicated conditions. **, p = 0.0073. Inset: Equal number of cells were lysed for western blot analysis.

(D) Relative *CXCL8* mRNA levels in HCT116 cells with stable knockdown of MLKL (shMLKL) or mismatch control (shMM) pretreated with combinations of 2 μM CpA, 20 μM zVAD and 1 μM NSA as indicated, and then stimulated with 0 or 2 ng/ml TNF for 3 h. Data is presented as mean with S.E.M. (n = 3). An unpaired t-test was used to test for the statistical difference between indicated conditions. *, p = 0.0108. Inset: Equal number of cells were lysed for western blot analysis. Line indicates that image was cut and spliced to remove non-relevant lanes from the scanned blot.

### Figure S7

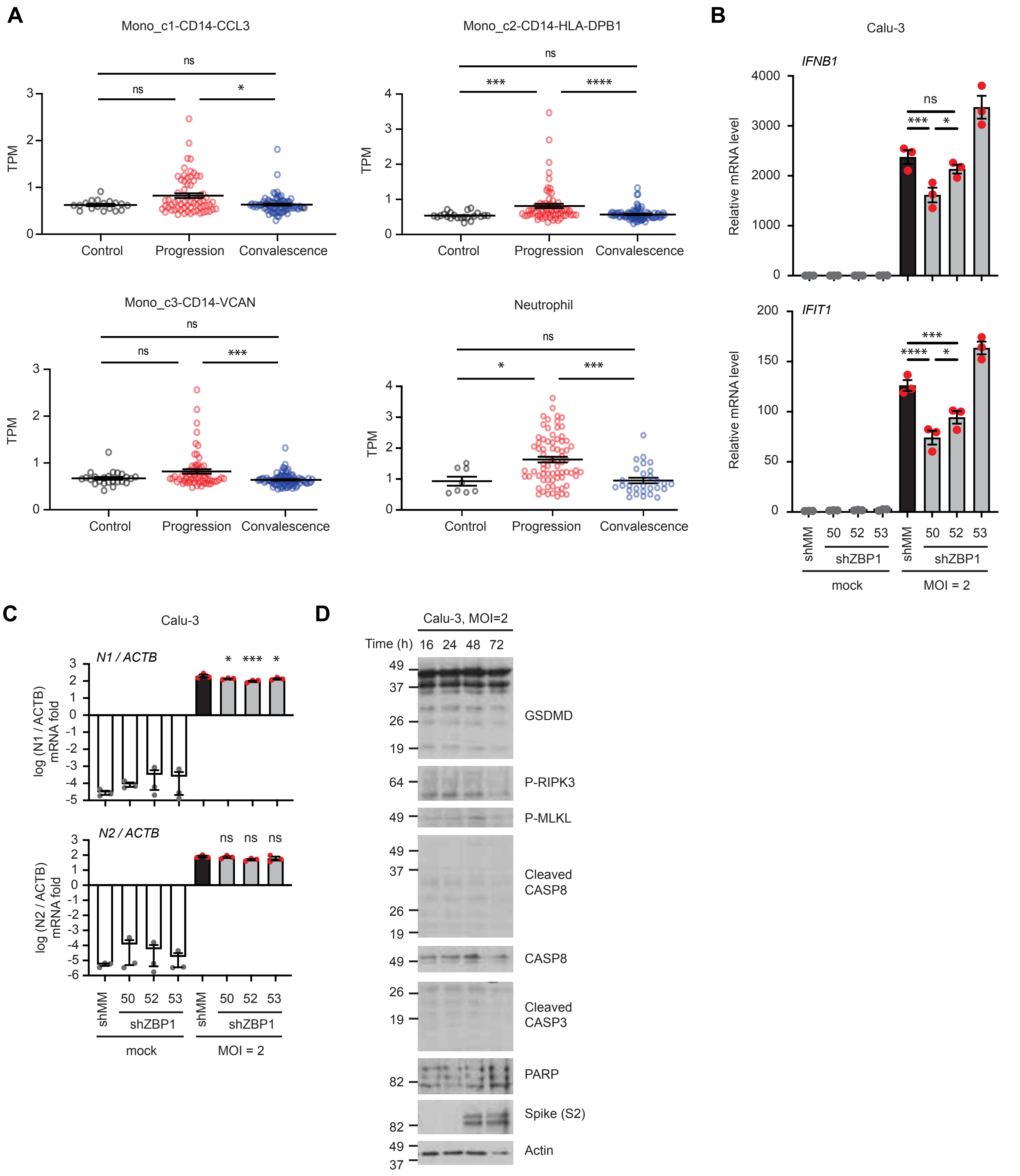

A
Mono_c1-CD14-CCL3
Mono_c2-CD14-HLA-DPB1
Mono_c3-CD14-VCAN
Neutrophil
TPM
Control
Progression
Convalescence
ns
*
***
****
B
Calu-3
IFNB1
IFIT1
Relative mRNA level
shMM
50
52
53
shZBP1
mock
MOI = 2
C
Calu-3
N1 / ACTB
N2 / ACTB
log (N1 / ACTB) mRNA fold
log (N2 / ACTB) mRNA fold
D
Calu-3, MOI=2
Time (h) 16 24 48 72
GSDMD
P-RIPK3
P-MLKL
Cleaved CASP8
CASP8
Cleaved CASP3
PARP
Spike (S2)
Actin

**Figure S7. Related to Figure 7. ZBP1 mediates SARS-CoV-2-induced inflammation**

(A) Patient-averaged single cell TPM values of *ZBP1* in subsets of peripheral blood cells of COVID-19 patients in progressive or convalescent stage, compared to healthy controls (Ren *et al.*, 2021). Data is plotted for individual patients with mean and S.E.M. Data is plotted as individual values with mean and S.E.M. Mono_c1-CD14-CCL3: n = 18 for control, n = 65 for progression, n = 81 for convalescence; Mono_c2-CD14-HLA-DPB1: n = 22 for control, n = 68 for progression, n = 88 for convalescence; Mono_c3-CD14-VCAN: n = 24 for control, n = 66 for progression, n = 89 for convalescence; Neutrophil: n = 8 for control, n = 73 for progression, n = 28 for convalescence. Kruskai-Wallis test and Dunn's multiple comparisons test were used to test for statistical differences between indicated conditions. ns, p > 0.3, *, p = 0.0305 in the mono_c1 panel, p = 0.0485 in the neutrophil panel, p<0.05, ***, p =.0.0004 in the mono_c2 panel, p = 0.0003 in the mono_c3 panel, p = 0.0001 in the neutrophil panel, ****, p<0.0001.

(B, C) Relative mRNA levels of *IFNB1* and *IFIT1* (B) or virus-encoded *N1* and *N2* (C) over β-Actin (ACTB, control) in Calu-3 cells with stable knockdown of *ZBP1* (shZBP1-50, shZBP1-52 and shZBP1-53) and control cells (shMM) infected with mock or SARS-CoV-2 virus at MOI = 2 for 72 h. Data is presented as mean with S.E.M (n = 3). (B) One-way ANOVA and Sidak's multiple comparisons test were used to test for statistical differences between indicated conditions. ns, p = 0.3831, ***, p = 0.0006 for *IFNB1*, p = 0.0003 for *IFIT1*, *, p = 0.0143, ****, p<0.0001. (C) One-way ANOVA and Sidak's multiple comparisons test were used to test for statistical differences between shMM MOI=2 and indicated conditions. *, p = 0.0294 for shZBP1-50, p = 0.0259 for shZBP1-53, ***, p = 0.0008, ns, p = 0.9359 for shZBP1-50, p = 0.0735 for shZBP1-52, p = 0.3109 for shZBP1-53.

(D) Western blot analyses of SARS-CoV-2-infected Calu-3 cells at MOI=2 for the indicated time.

#### Supplementary material and methods

##### *Cell culture*

U2OS / NOD2 cells (Fiil et al., 2013) were cultured in DMEM supplemented with 10% (v/v) FBS, 60 µg/ml penicillin and 100 µg/ml streptomycin. HL60 cells were cultured in RPMI media supplemented with GlutaMAX, 10% (v/v) FBS, 60 µg/ml penicillin and 100 µg/ml streptomycin (PS). Differentiated HL60 cells were obtained by culturing in complete growth media supplemented with 1.3 %DMSO (v/v) (Sigma D2650) for 7 days in culture.

HEK293FT / RIPK3-2xFV cells were generated following same protocol as described in the main text. After selection, cells were subject to FACS sorting giving rise to a low-mCherry RIPK3-expressing HEK293FT / RIPK3-2xFV population.

##### *Generation of MLKL stable knockdown cell lines*

HCT116 / RIPK3-2xFV cells knockdown for MLKL were generated same as described in the main text using shRNA with the following sequences:
shMLKL-A, SHCLND-NM_152649, TRCN0000196741,
CCGGGAGTCAAATCTACAGCATATCCTCGAGGATATGCTGTAGATTTGACTCTTTTTTG;
shMLKL-B, SHCLND-NM_152649, TRCN0000194846,
CCGGCCTCTGACAGTAACTTTGATACTCGAGTATCAAAGTTACTGTCAGAGGTTTTTTG.

##### *Generation of U2OS / NOD2 cells stably expressing RIPK3 variants*

For the stable expression of RIPK3 and its derivatives in U2OS / NOD2 cells , RIPK3, RIPK3-2xFV and RIPK3dC-2xFV were subcloned into pBABE-puro plasmids using primers 5’-ACGCGTATGTCGTGCGTCAAGTTATG-3’ and 5’-GTCGACTTACTTATCGTCGTCATCCTTGTAATCTTTCCCGCTATGATTATACCAAC-3’ (matching the C-terminus of RIPK3, adding FLAG tag sequence) or 5’-GTCGACTTACTTATCGTCGTCATCCTTGTAATCTTCCAGTTTTAGAAGCTCCAC-3’ (matching the C-terminus of FV, adding FLAG tag sequence) to amplify the corresponding sequences from LZRS-zeo plasmids.

U2OS / NOD2 cells were transduced with retroviral particles from the above plasmids and selected with puromycin as described in the main text.

***qPCR primers used in supplementary figures***

*A20* (*TNFAIP3*): 5'-ATGCACCGATACACACTGGA-3' and 5'-GGATGATCTCCCGAAACTGA-3'.

*IFNB1*: 5'-ATGACCAACAAGTGTCTCCTCC-3' and 5'-GGAATCCAAGCAAGTTGTAGCTC-3';

*IFIT1*: 5'- GCGCTGGGTATGCGATCTC-3' and 5'- CAGCCTGCCTTAGGGGAAG-3'.

***One-step qPCR***

One-step RNA-to-Ct qPCR was used to determine the amount of intracellular SARS-CoV-2 following manufacturer's protocol (Thermo Fisher 4392938). 100 ng RNA was loaded into each reaction with primers and probes for SARS-CoV-2 (IDT 10006713) or β-Actin (Thermo Fisher 4331182, Assay ID Hs00357333_g1) at manufacturer's recommended concentrations.

***Site-directed mutagenesis***

RIPK3 mutants used in this study (K50A, D142N, RHIMmut) were generated using a Q5 site-directed mutagenesis kit (NEB) on the LZRS-zeo-RIPK3-2xFV plasmid according to the manufacturer's protocol. The following primers were used for mutagenesis:

K50A, 5'-TGTGGCGGTCGCGATCGTAAACTCG-3' and 5'-TCGTAGCCCCACTTCCTA-3';

D142N, 5'- CCTGCACCGGAACCTCAAGCC-3' and 5'-AGCACCGGGTTCTGGTCG-3',

RHIMmut, 5'-GCTGCAGGAGACAACAACTACTTG-3' and 5'-TGCCGCCCCAGAGCAGTTGTATATG-3'.

##### *Additional antibodies used in supplementary figures*

Anti-phospho-IκBα (Cell Signaling 2859), anti-IκBα (Cell Signaling 9242), anti-phospho-ERK1/2 (Cell Signaling 4370), anti-ERK1/2 (Cell Signaling 4695), anti-2A peptide (Millipore ABS31), anti-XIAP (BD 610762), anti-Gasdermin D (GSDMD, Cell Signaling 97558), anti-phospho-MLKL (Cell Signaling 91689), anti-MLKL (Cell Signaling 14993), anti-PARP (Cell Signaling 9542), anti-SARS-CoV-2 Spike protein (GeneTex GTX632604).

##### *Chemokine array*

The chemokine array was performed following the manufacturer's protocol of Proteome Profiler Human Chemokine Array Kit (R&D ARY017). Pixel intensity from each chemokine dot area were quantified using Fiji. Of each detectable chemokine, fold change of Dox-treated condition over DMSO-treated condition was divided by the fold change of pixel intensities of positive control dots for normalization.

##### *Cytokine array*

The cytokine array was performed following the manufacturer's protocol of Proteome Profiler Human XL Cytokine Array Kit (R&D ARY022B). Individual mean pixel intensity of each dot for each exposure time was quantified using Python. Raw pixel intensities were inverted (subtraction from 255 for 8-bit image) and background normalized by subtracting the mean intensity of each blot for fold-change analysis. The fold change of Dox-treated condition over DMSO-treated condition was normalized by dividing by the fold change of pixel intensities of positive control dots. The normalized fold change at different exposure times were plotted asa heatmap.
